# Age and sex alter the immune response in a chronic fibrosis model via changes in T cell and macrophage phenotype

**DOI:** 10.64898/2026.08.27.747581

**Authors:** Joscelyn C. Mejias, Anna Ruta, A Shri Ramanujam, Katlin B. Stivers, Sean Kelly, Natalie Rutkowski, Kavita Krishnan, Locke Davenport Huyer, Chris Cherry, Franck Housseau, Savannah Est-Witte, Jennifer H. Elisseeff

**Author notes:** Current institution (Joscelyn C. Mejias, Locke Davenport Huyer, Franck Housseau).

## Abstract

The foreign body response (FBR) is an immune mediated event that occurs with every material implant. The extent of the fibrosis is dependent on many factors including the biomaterial design, tissue location, and host factors such as age, sex, ancestry, diet. There are known clinical outcomes of implants dependent on age and sex, including increased fibrosis and implant failure in aged and female patients. As the population ages, there is a growing need to understand how aging affects the FBR, and how preclinical models can capture this to guide biomaterial design. Here, we investigated how chronic fibrosis in a murine model of the FBR is altered by two biological factors: age and sex. We investigated changes in fibrosis using a volumetric muscle loss (VML) injury model coupled with polycaprolactone (PCL) or polyethylene (PE) microparticle implants. Fibrosis was quantified through gene expression, microscopic analysis of histologic sections, and the corresponding immune response measured via gene expression and flow cytometry data. We found gene expression differences with immune pathways enriched in female mice, and microscopy revealed collagen birefringence area increased in young male mice. Both the innate and adaptive immune response were altered by age and sex via T cell and macrophage phenotype, and the effects of aging differed between sexes. These results reveal both variables contribute to discrepant outcomes in both fibrosis and the local immune response to synthetic material implants. This demonstrates a clear need to understand and account for the influence of biological factors in biomaterial design.

## Introduction

The global population is aging, with the World Health Organization estimating 1 in 6 people will be over the age of 60 by 2030[1]. Older patients have disproportionately greater clinical need for biomedical implants and materials[2], and aging is also accompanied by biological changes that can alter the response to these devices. Likewise, patient sex has been identified as a key factor in clinical success of biomedical implants and materials, with female patients more likely to experience adverse events from medical devices[3–5] and implant materials[6–8] including higher rates of allergic skin and tissue reactions, aseptic loosening, and stronger inflammatory responses to foreign materials[9]. Together, an aging population and the recognition of sex and age as important clinical variables underscore the need to understand how host biology shapes responses to preclinical biomaterials, and ultimately, clinical implant and material performance.

Biomaterials have applications in drug delivery, immunotherapy, and tissue regeneration[10–12]. Any material implant will initiate the immune-mediated foreign body response (FBR), resulting in a fibrous encapsulation of the implant. The FBR is a complex process initiated by protein adsorption onto the material surface, which drives complement activation leading to the recruitment of neutrophils, monocytes, and macrophages[13]. As the inflammatory response proceeds from acute to chronic, macrophages fuse together into foreign body giant cells, the adaptive immune system infiltrates, and both work in concert with local stromal cells to create a fibrotic capsule that isolates the implant from the surrounding tissue[14–18]. This often leads to implant failure and negative outcomes for the host[19]. The extent of the FBR is dependent on several factors, some inherent to the biomaterial (i.e. size, composition, topography)[15, 20–22] and some inherent to the host (e.g., age, sex, disease history, tissue locale)[19, 23]. The field of biomaterials-based tissue regeneration focuses on understanding material properties and modifications that can minimize fibrosis or leverage wound healing response for improved tissue integration, but there is a critical need to consider host factors that will alter the immune response, and thus, a material’s therapeutic efficacy. Here we focus on how age and sex affect the fibrotic response to implanted biomaterials in mice.

Aging is associated with a general reduction in immune function that leads to greater susceptibility to disease, hindered response to infection, and lowered regenerative capacity[24, 25]. The aged adaptive immune system shows decreased T cell diversity, reduced CD4+ and CD8+ T cell proportions, and diminished B cell function, while the aged innate immune system exhibits increased neutrophils and monocytes but with impaired phagocytic capabilities[24–26]. Macrophage polarization skews toward pro-fibrotic phenotypes with age[26, 27]. The immune response shifts from type-2 dominance with elevated interleukin (IL)-4 and IL-13 in young mice towards TNF-α and IL-1β-driven chronic inflammation in aged mice[28]. Aging also promotes fibrosis pathways over regenerative ones across multiple organs including lung, liver, and kidney[29], which is recapitulated in several human disease contexts, where aged patients have higher fibrosis scores than their younger counterparts[29–33]. This shift is attributed both to immune dysfunction and an accumulation of senescent cells, which produce pro-fibrotic mediators[34, 35] as part of their senescence-associated secretory phenotype (SASP). It follows that the FBR is altered by aging, though the effect depends on implant material; Some studies report aged mice show delayed resolution of inflammation following implantation, with sustained pro-inflammatory macrophage responses and impaired M2-like polarization compared to young mice, though the opposite effect has also been observed[27, 36].

Sex also influences in the immune system and FBR, mediated by gonadal hormones (e.g., testosterone, estrogen, progesterone) and the chromosomal complement, which is the complete set of autosomes and sex chromosomes present in a cell. Murine transcriptomic analysis demonstrated that sexual dimorphism in immune cells is most pronounced in macrophages from different tissues, with male macrophages and neutrophils expressing more pro-inflammatory markers (TNF-α, IL-1β) and TLRs[37–40]. In a bleomycin-induced model of pulmonary fibrosis, greater upregulation of pro-inflammatory and extracellular matrix (ECM) genes expression by myeloid cells in males is associated with the significantly increased development of lung injury, inflammation, and fibrosis compared to females[38]. However, one study with polydimethylsiloxane implantation showed that female Balb/c mice exhibit significantly higher IL-1β expression [41]. These studies highlight the context-dependent nature of studying sex differences in immunity, underscoring the need to characterize them specifically in the context of the FBR.

Here we evaluate the age- and sex-specific differences that occur in an established murine model of traumatic muscle injury and material-induced fibrosis[42] at both the transcriptional and protein levels across the stromal and immune microenvironment. We demonstrate that these biological factors influence the balance of innate and adaptive immune cells as well as local fibrotic factors and must be accounted for in bioengineering approaches to reverse or prevent pathological fibrosis.

## Results

### Inflammatory and ECM-associated genes are upregulated in female mice, regardless of age in a biomaterial-induced chronic fibrosis model

To investigate the effects of age and sex on the chronic fibrosis associated with the FBR, we implanted fibrosis-inducing polycaprolactone particulate (PCL, MW ~50,000 Da) into a VML injury model in C57BL6 male and female mice at 2 months (young) and 18 months (aged) (Fig. 1A). After 6-weeks, the injured muscle along with fibrosis-encapsulated PCL implant were excised and bulk RNA sequencing (bulkseq) was performed. As differential gene expression analysis of bulkseq data is conducted pairwise, we ran 4 independent comparisons: young female (YF, n = 5) vs young male (YM, n = 5), aged female (AF, n = 5) vs aged male (AM, n = 5), young female vs aged female, and young male vs aged male (visualization of intersection and differences between each comparison is in (Fig. S1A). Fewer significant differentially expressed genes were observed when comparing between age groups within the same sex (< 200) than when comparing genes between sexes within the same age group (> 3,000) (Fig. 1) and S.File 1). Gene set enrichment analysis (GSEA) using Hallmark[43–45] and KEGG[46–48] was used to identify pathways enriched in male or female mice within each age group. Adipogenesis and myogenesis gene sets were enriched in both young and aged male mice relative to their age matched female counterparts while inflammatory response and interferon pathways were enriched in females relative to the males; immune-related pathways are highlighted in green (Fig. 1D and S1B–C). GSEA by age (within each sex) showed fewer significant pathways and few unique leading-edge genes, likely due to the low number of differentially expressed genes between age groups of the same sex. We then ran GSEA with the Matrisome Project Database[49] gene sets and found that ECM-affiliated, regulators, glycoproteins, collagens, and core matrisome pathways were enriched in the female samples relative to age-matched males (Fig. S2). When comparing by age within each sex the matrisome pathways are primarily enriched in young, but the leading-edge genes are again common between the sets.

**Fig. 1.**
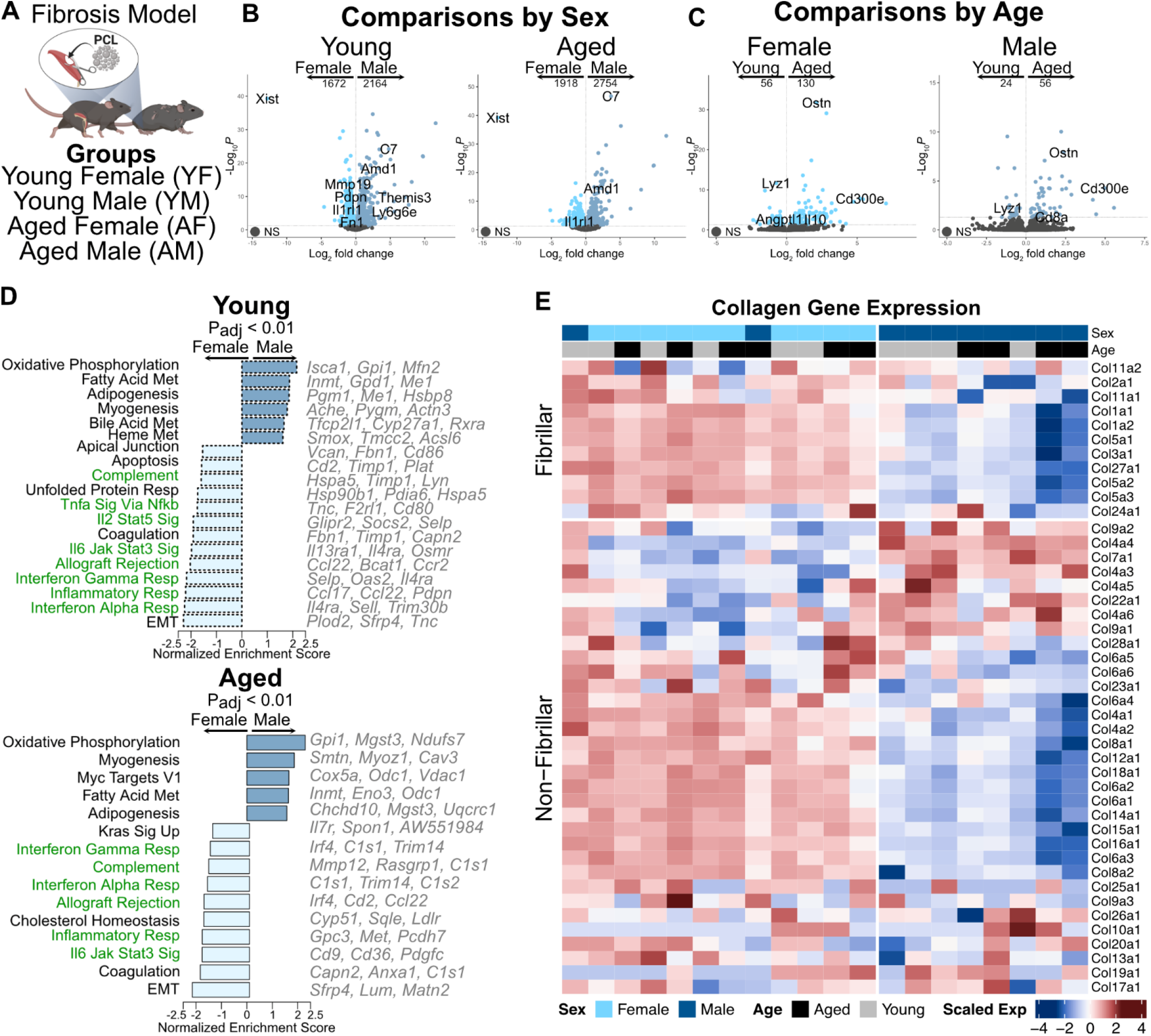
The chronic FBR is altered by sex and age. (A) Schematic representation of injury and groups analyzed including young (2 month) and aged (18 month), male and female C57BL6 mice received a VML injury treated with PCL that was excised 6 weeks later. (B-C) Volcano plots for each comparison set by either age or sex with a select set of genes shown across each plot. (D) Hallmark GSEA (Padj < 0.01) of the sex comparisons within each age group (young or aged) shows distinct pathways up in male vs female at both ages. Gene sets related to the immune system are highlighted in green. (E) Hierarchical clustering of the collagens within the dataset split by fibrillar vs non-fibrillar with scaled expression by row.

As collagens are critical to fibrosis[50], we next examined the different collagen isoforms captured within the dataset. Row-scaled gene expression was visualized by heatmap, with collagen genes grouped into fibrillar and non-fibrillar and hierarchically clustered within groups. Unsupervised hierarchical clustering of samples separated them into two groups that closely corresponded to sex (Fig. 1E). Regardless of age, female mice showed elevated expression of fibrillar, and to a lesser extent non-fibrillar, collagens, suggesting sex-specific matrix accumulation and remodeling.

### Age, sex, and implant type affect expression and distribution of proteins contributing to fibrosis

To collect histological information on the changes in fibrosis, including protein level changes and spatial information, we analyzed Masson’s Trichrome, which highlights collagens, and Picrosirius Red (PSR), which measures the birefringence of collagens under polarized light, an established method of quantifying fibrosis, scar formation, and collagen organization in injured or pathological tissues[51, 52], on 6-week VML PCL implants. For these experiments we included a second particulate material, polyethylene (PE, UHMW, 125 µm average size), to investigate whether implant type plays a role in age and sex FBR differences. By 6-weeks post implantation, both materials induced the FBR and developed substantial fibrosis, however the PCL groups displayed higher percentages of percent FBR Collagen and percent Red PSR, indicating thicker, more mature collagen fibers than the PE groups (Fig. S3A and 2A–D). QuPath was used to quantify the blue collagen staining in Masson’s Trichrome within the fibrosis area (Fig. 2A) and showed no significant differences due to age or sex within the PCL groups. However, within the PE groups aged female mice have a significant increase in collagen percentage compared to age-matched males and young females (Fig. S3B and 2B). Further characterization by PSR (Fig. 2C and S4) found an increased percent red in young males for both PCL and PE groups, indicating higher red birefringent collagen signal, suggesting thicker or more mature collagen fibers (Fig. 2D). The ratio of red to green birefringence was not significantly different between sex or age within either implant group (Fig. S4B).

**Fig. 2.**
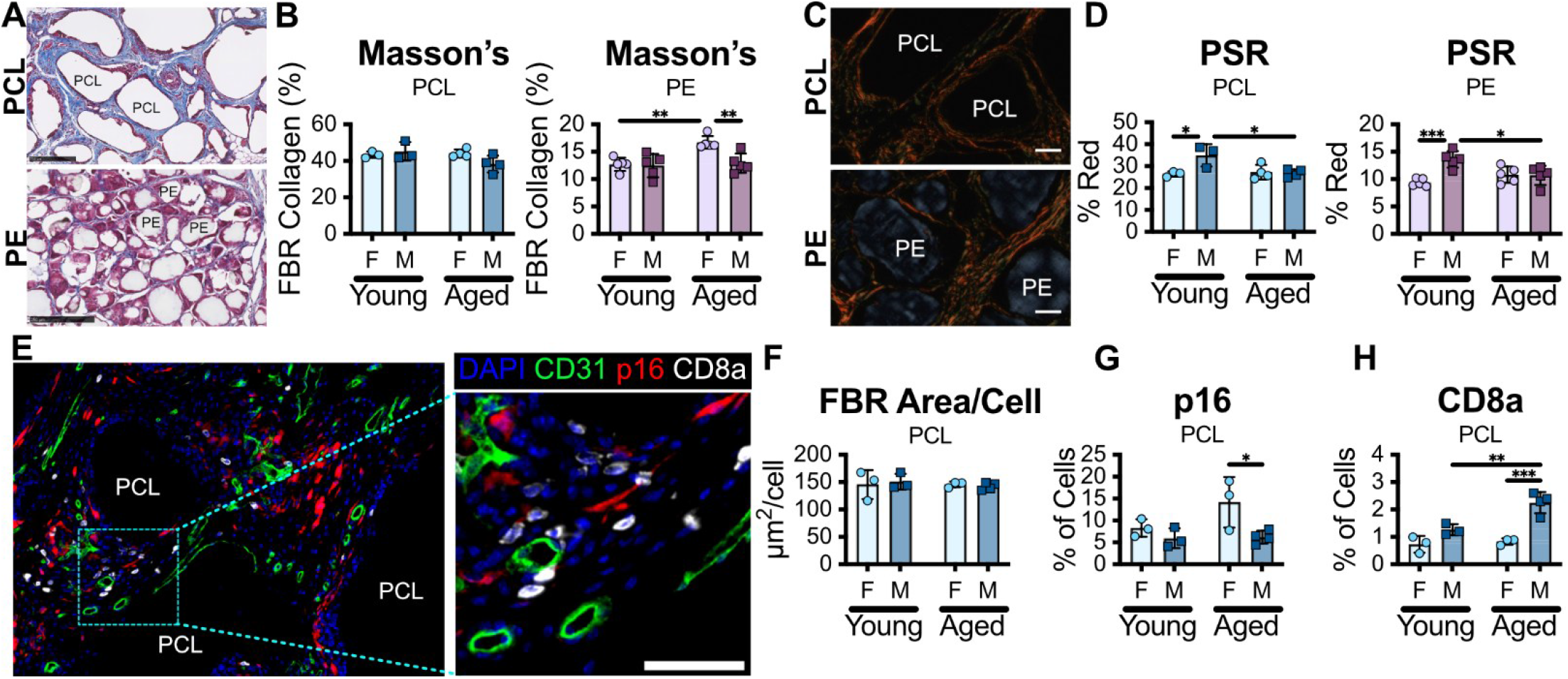
Fibrosis quantification is altered by age, sex, and material. (A) Representative images of Masson’s trichrome for PCL (plot in blues) and PE (plot in purples) 6 weeks after VML, scale bars = 250 µm (B) Quantification of the collagen (blue) area within the FBR shows unique increase in the PE aged female. (C) Representative Picrosirius Red (PSR) imaging, scale bars = 50 µm (D) Quantification of the red channel shows both materials are significantly higher in the young males. (E) Representative immunofluorescence staining (aged female) of vessels (CD31), senescence (p16) and CD8 T cells (CD8a), scale bar = 50 µm, images linearly contrasted for clarity. (F) Quantification of the number of cells per area. (G) p16 a marker for cellular senescence is significantly higher in aged female than male. (H) CD8a is significantly higher in aged males compared to aged females and young males. Statistics: Two way ANOVA with Tukey post-hoc analysis between YF-YM, AF-AM, YF-AF, YM-AM.

Within the bulkseq dataset, we observed both stromal and immune gene set differences between age and sex groups that could contribute to expressed genes within at least one of the four comparisons, (Fig. 1B–D). CD8 is a marker for cytotoxic T cells, one of the major cell populations that contributes to IFNγ production, while IL-6 is a cytokine associated with the SASP of senescent cells, which accumulate with aging and fibrosis and are commonly identified by their expression of p16. To investigate these differences on the protein level, we co-stained these samples for CD31 (vessels), p16 (senescent cells), and CD8 (T cells) (Fig. 2E) shows a representative example of immunofluorescence (IF). Since the PE particulates exhibited autofluorescence that interfered with IF, we only analyzed the PCL implant. Within each FBR region, the area analyzed was normalized to the number of nuclei within the region of interest (ROI) to ensure group differences were not driven by variations in cell numbers (Fig. 2F). There was no difference in the number of vessels or their distribution by size (Fig. S5B–C). However, there was a sex-specific difference in the percentage of p16+ cells in the aged samples (Fig. 2G), and an age and sex specific increase in CD8+ T cells in the aged male condition, consistent with the increased expression of C*d*8a in aged male mice compared to young mice, (Fig. 2H and 1C). These results represent a complex interplay of age and sex as variables impacting chronic fibrosis in the muscle. Taken together with the bulkseq dataset for PCL implants, the sex-biased differences in inflammatory gene expression and fibrillar collagen do not result in significantly more organized or thicker collagen structure in female mice, as would be suggested by gene expression alone.

### Age and sex alter T cell phenotype in biomaterial-induced fibrosis by shifting CD4/CD8 ratios toward CD4 in female mice and CD8 in aged mice

T cells are critical to the development and maintenance of fibrosis. Both CD4+ and CD8+ T cells can regulate fibroblast and macrophage function. To better understand T cell dynamics in chronic fibrosis we visually confirmed both CD8+ and CD4+ cell types within the fibrosis encapsulating the male PCL implants with IF staining (Fig. 3A and S6). Using flow cytometry, we investigated the chronic immune microenvironment in 6-week VML Saline, VML PCL, and VML PE tissues (S.Table 1 and Fig. S7). Of note, there were some differences in quadricep muscle weight based on sex with and without the particulate, (Fig. S8). Proportion and cell count data is reported with respect to the number of whole quadriceps collected for analysis regardless of injury or material, as isolation of the fibrotic region alone was not feasible. We found the number of total CD3+ T cells significantly increased in the aged saline female mice, however there were no changes in CD3+ T cells counts associated with age or sex within the PCL and PE conditions (Fig. 3B). Across all VML injury conditions, the proportion of CD8+ T cells increased with age, with a sex-specific increase in aged males compared to aged females (Fig. 3C–E and S6B; Fig. S9A). In the PE and PCL conditions, this increase in CD8+ T cell frequency also correlated with an increase in CD8+ T cell count (Fig. S9B). In both the PCL and PE biomaterial-induced fibrosis conditions, the proportion of CD4+ T cells increased in the female, aged group, and in both female groups in the PCL condition (Fig. 3D–E).

**Fig. 3.**
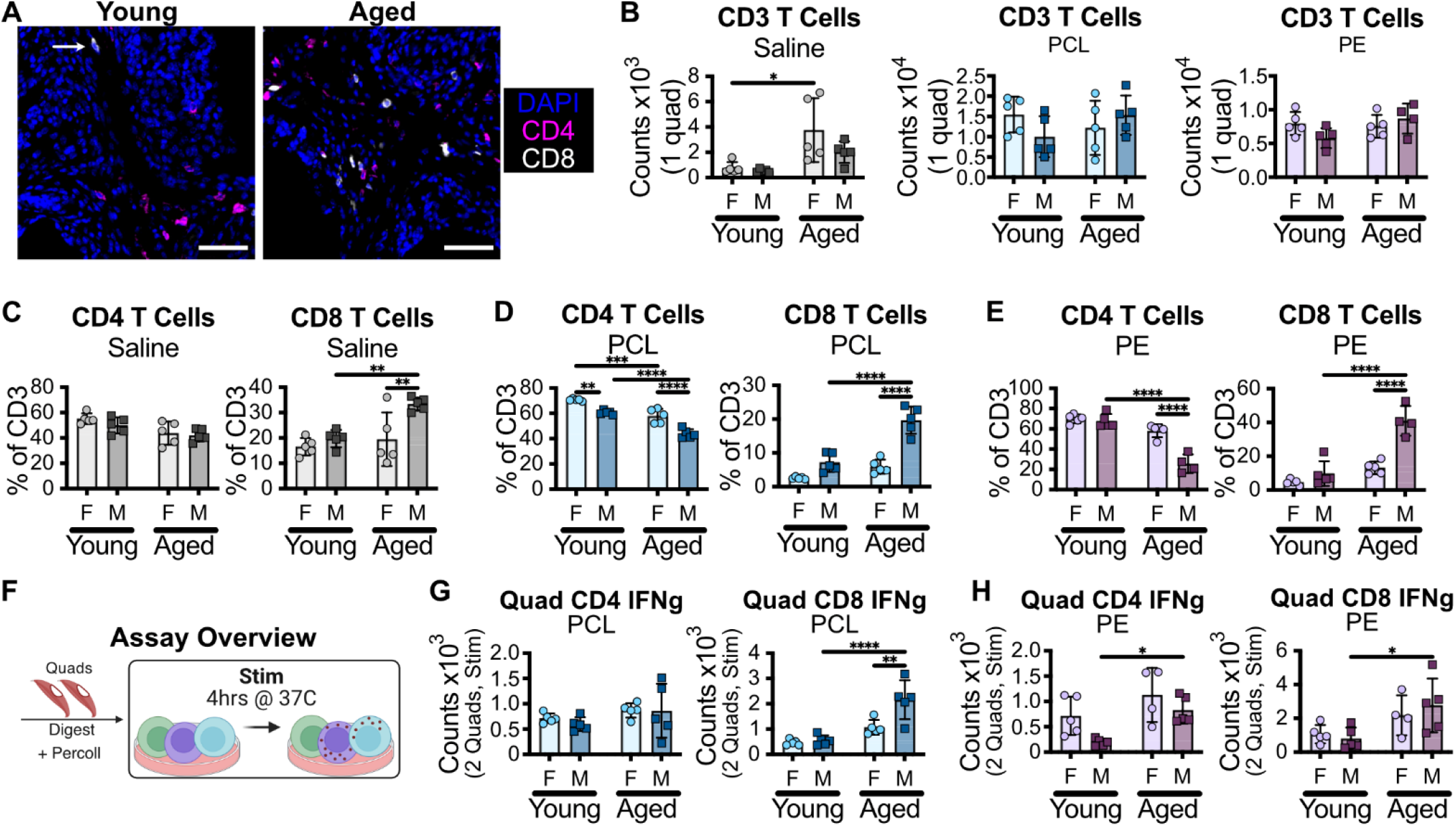
Age and sex alter T cell phenotype. (A) Representative fluorescent images of nuclei (DAPI), CD4 and CD8 in the 6 week VML PCL induced fibrosis, scale bars = 50 µm (B) CD3+ T cell counts of quadriceps from 6 week VML-Saline (greys), -PCL (blues), and -PE (purples). Proportions of CD4+ and CD8+ within CD3+ T cells for (C) Saline (D) PCL and (E) PE. (F) Schematic overview of cell stimulation protocol. Ifng counts for CD4+ and CD8+ T cells in the quadriceps treated with (G) PCL and (H) PE. Statistics: 2 way ANOVA with Sidak posthoc analysis comparing only YF-YM, YF-AF, YM-AM, AF-AM, adjusted p = *0.05, **0.01, ***0.001, ****<0.0001.

CD4+ and CD8+ T cell effector phenotype and function can be described through the cytokines they produce; CD4+ effector subsets include IFNγ-producing Th1, IL-13-producing Th2, and IL-17-producing Th17. To understand the phenotypic potential of CD4+ and CD8+ T cells, cells from both inguinal lymph nodes (iLNs) and quadriceps with FBR were stimulated *ex vivo* for 4 hours then stained with an intracellular cytokine staining (ICS) panel for flow cytometry analysis (Fig. 3F, S.Table 2, and Fig. S10). For quadriceps, since non-material conditions have low numbers of CD3+ T cells, data was only collected from 6-week VML PCL and VML PE conditions.

The total number of CD3+ T cells in the iLNs declined with age regardless of sex with a 10-fold decrease for all conditions (10^6^ in young mice to 10^5^ in aged mice) (Fig. S11); however, within the quadriceps, the CD3+ T cell counts were not significantly different across age and sex within implant conditions (Fig. S13). All three characteristic cytokines (IFNγ, IL-13, and IL-17) were expressed by CD4+ T cells within the quadriceps regardless of sex or age, with IFNγ as the dominant cytokine in all groups except the female PCL mice in both age groups (Fig. S12). In the iLNs, all three cytokines were only significantly expressed in the aged groups, with a higher proportion in female vs male aged mice (Fig. S12–13). IFNγ was also the dominant cytokine in all significant iLN groups (aged) and is a major contributor to fibrosis through both CD4+ Th1 and CD8+ T cells. Within the fibrotic quadriceps, both CD4+IFNγ+ and CD8+ IFNγ+ significantly increased in aged male mice across both material types, except for the CD4+ IFNγ+ PCL, reflecting a consistent sex difference within the aged group (Fig. 3G and S13). For IFNγ, there were significant differences in cytokine expression within the fibrotic quadriceps that were independent of the changes within the iLN. Tissue and condition specific cytokine changes were also observed for IL-17, IL-13, IL-2 and IL-10, each with different trends by age, sex, and tissue dependency, (Fig. S15). This emphasizes the need to assess regional tissue responses along with the host demographics when evaluating biomaterial responses.

### Chronic fibrosis from synthetic implants skews muscle macrophages in female mice toward an M1 phenotype regardless of age

Myeloid cells and in particular macrophages, make up the largest proportion of immune cells within the FBR. IF imaging for the myeloid lineage marker CD11b showed myeloid cells distributed throughout the fibrosis with a high localization around the empty PCL particulate space (Fig. 4A and S16). UMAP of down sampled flow cytometry visualized the distribution of major cell types, while condition-specific density plots highlighted changes in relative population abundance across each sample type (Fig. 4B, S.Table 1, and Fig. S7). The total CD45+ immune cell counts were not different by age or sex although the number of total immune cells significantly increased with material implants (Fig. S17A). The largest immune cell population was F4/80+ macrophages which, as a proportion, of total immune cells had a slight decline for the PCL males with age (Fig. S17B). Using flow cytometry, we identified 6 other myeloid populations (e.g., eosinophils, neutrophils, monocytes, DCs) as well as lymphoid B and NK cells; each condition had some minor shifts in proportions by age and sex but no shared changes across the material conditions (Fig. S17).

**Fig. 4.**
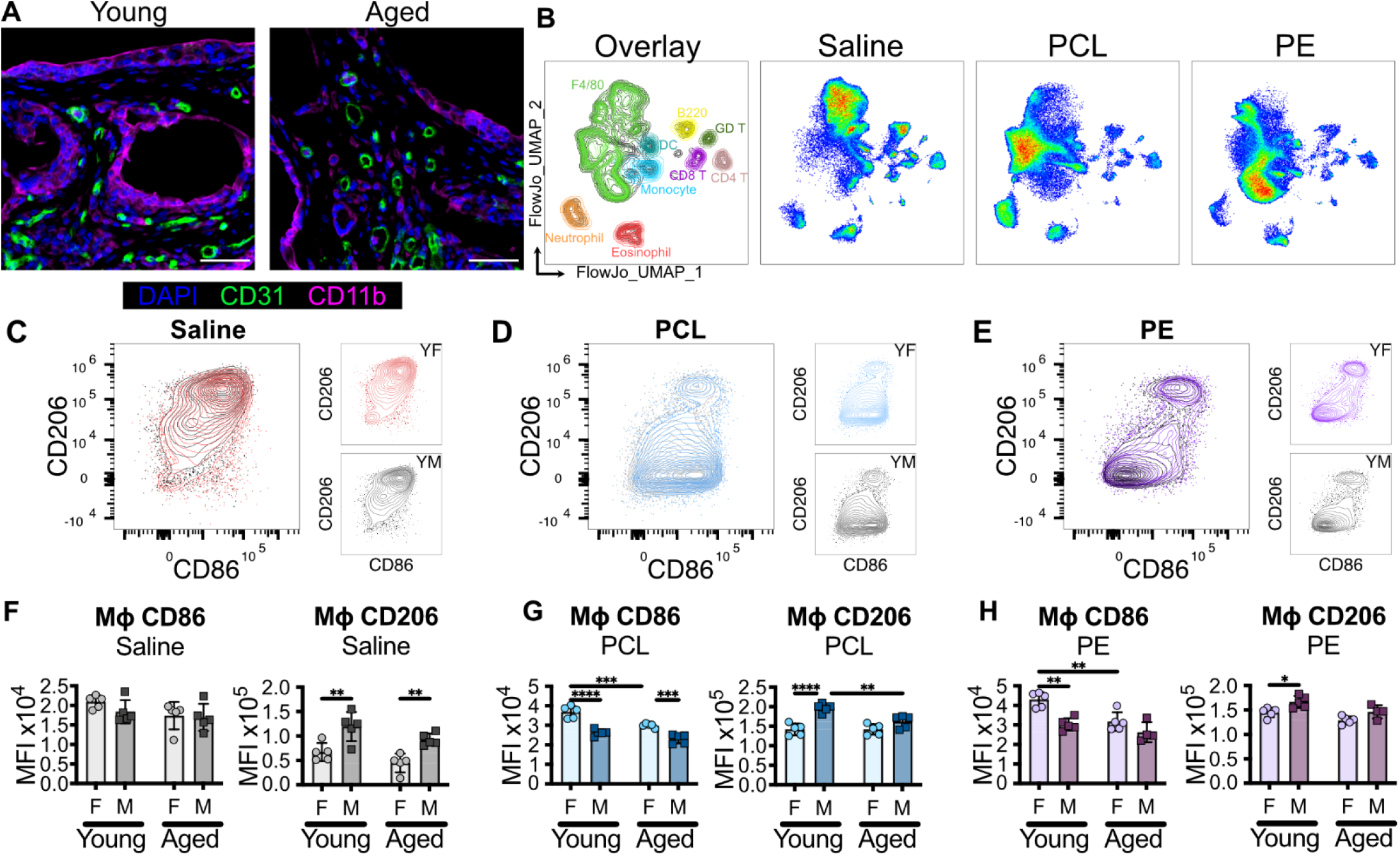
Macrophage phenotype is altered by age, sex, and material. (A) Representative fluorescent images of nuclei (DAPI), vessels (CD31) and myeloid cells (CD11b) in the PCL induced fibrosis; images linearly contrasted for clarity, scale bars = 50 µm (B) UMAP projection of all conditions with downsampled individual samples with gated populations labeled in the overlay and split by individual condition. (C) Cell counts for immune cells (D) F4/80 Macrophage counts within each condition. (E-G) CD86 and CD206 median fluorescence intensity from the F4/80 Macrophage gate for Saline, PCL and PE. Statistics: 2 way ANOVA with Sidak posthoc analysis comparing only YF-YM, YF-AF, YM-AM, AF-AM, adjusted p = *0.05, **0.01, ***0.001, ****<0.0001.

Cell surface markers CD86, CD206, and MHCII are key to the function of macrophages. CD86 is the ligand for T cell activation through CD28 and CTLA4, CD206 is a mannose receptor implicated in ECM rearrangement, and MHCII is primarily associated with CD4+ T cell activation. In the injury only condition (VML Saline), median fluorescence intensity (MFI) of these markers on macrophages showed a sex-specific increase in CD206 expression on male-derived macrophages regardless of age and no difference in CD86 expression (Fig. 4C and 4F); however, with the presence of PCL and PE material implants, the sex difference in CD206 expression was lost in aged, and the young female mice had significantly increased CD86 expression compared to the young male and aged females (Fig. 4C–H). MHCII expression had a minor decrease in young males compared to young females for PE (Fig. S18A). While CD86 and CD206 are canonical markers used to differentiate M1-like proinflammatory and M2-like pro-regenerative macrophages respectively, the CD206+ population found in a 6-week chronic injury with significant fibrosis was most likely associated with fibrotic remodeling and not a regenerative response.

Gating of macrophage subpopulations further emphasized both age and sex dependent expression of CD206, CD86 and MHCII (Fig. S18B). The largest population for the injury only condition (VML Saline) was a subset that was double positive (CD206+CD86+), while the largest proportion of macrophages for the material--induced fibrosis were CD206 low/negative (Fig. S18C). *Mrc1* (CD206) and *Cd86* (CD86) double expression at the gene level was observed to be broadly expressed in naïve skeletal macrophages[53] and *Mrc1* (CD206) along with *Mgl2* and *Clecl10a* (murine CD301a and CD301b) at higher expression in naïve quadriceps and diaphragm macrophages relative to peritoneal and lung alveolar macrophages[54]. Within the largest material-related macrophage population (labeled Mac 1; CD206 low/negative), the CD86 expression was sex dependent in all injury conditions (Saline, PCL, and PE) and the MHCII expression was sex dependent in PE and both age and sex dependent in PCL (Fig. S18D). The CD206+CD86+ population was further defined by the expression of CD301b with the positive population labeled Mac 2 and the negative Mac 3 (Fig. S18B).

CD301b is a lectin with specificity for galactose (Gal) and n-acetylgalactosamine (GalNAc). There was a proportional decrease in the of Mac 2 (CD301b+) in the aged PCL that was not observed in Saline or PE (Fig. S19). While there were some condition-specific changes by age and sex across both Mac 2 and Mac 3, there was often an increased CD206 expression (measured by MFI) on male-derived macrophages. Specifically within the materials-induced fibrosis, there was an increased CD86 expression on the female-derived macrophages with an overall age-related decline in CD86 expression for both sexes (Fig. S19). While there were limited changes in the relative proportions of broad immune subtypes investigated, there was a clear phenotypic change associated with age and sex in macrophages, and the CD86 expression differences were only evident with implanted materials. As immunomodulatory materials are developed to promote tissue regeneration by rebalancing M1-like and M2-like expression, there will be limited translational efficacy if age and sex are not accounted for.

## Discussion

The growing use of biomaterials in aging patients and the recognized influence of sex on implant outcomes highlight the need to define how age- and sex-dependent host biology shapes FBR and material performance Chronic fibrosis by the FBR to a material implant is a dynamic immune response involving both the innate and adaptive immune arms communicating with the local tissue microenvironment. Biological host factors such as age, sex, infection history, and genetics affect immune cell function, and while it has been established that material properties (e.g., size[22], shape[22], and stiffness[55–57]) influence immune cell responseto implanted biomaterials, there are still significant gaps in the intersection of how biological factors inherently control and influence these material-associated immune responses.

Our findings show that age and sex can alter the stromal, innate, and adaptive immune response in multiple fibrotic microenvironments. We found that age drove a significant shift in CD4+ to CD8+ T cell distribution, and sex drove a significant shift from CD8+ to CD4+ T cell dominance in female mice (PCL, PE aged), which has also been seen in mouse studies of adipose by age and sex[58] and naturally derived material implants by age[42] and sex[59]. We also identified canonical cytokines for most major T helper cell phenotypes (Th1, Th2, Th17). As T cell phenotypes are heavily influenced by the myeloid response, we also found significant heterogeneity in the macrophage response.

Macrophages are the largest immune cell population by both count and proportion within both material fibrosis models. They are often the primary target in immunomodulatory materials to balance an M1/M2-like phenotype to promote tissue regeneration, although the M1/M2 paradigm is increasingly recognized as a simple representation of a much more complex phenotypic space[60–62]. The VML itself is a chronic, non-healing wound environment, and while there are sex differences in CD206 expression and shifts in CD86 expression observed by age, the specific CD86 expression increases in females are only apparent in the material-induced fibrosis. Methods targeting the balance of CD206/CD86 may not be optimal for all populations if only tested in one sex or one age group. This highlights the need of biomaterial work to address the mandates of testing in both sexes and at least consider the consequences of aging to ensure that they modulate the local immune microenvironment as intended.

We also observed differences between data modalities. At the transcriptional level within the PCL fibrosis, we observed significant differences in gene set enrichment between sexes, while at the protein level, age became an additional driver of variability. At the transcriptional level, inflammatory and ECM genes including both fibrillar and non-fibrillar collagen associated genes were primarily upregulated in female mice, regardless of age. This would suggest an increased fibrotic response in female mice, but this was not observed by immunofluorescence, Masson’s trichrome staining, or Picrosirius Red imaging in the corresponding fibrosis quantification (PCL). Importantly, the baseline age-associated shifts observed in naive mice did not predict the local chronic FBR for either implant form factor. Taken together, this data suggests that predicting therapeutic outcomes requires context-specific evaluation rather than relying on baseline data, reinforcing the need for multi-targeted strategies rather than single-pathway interventions.

As the field of tissue regeneration seeks to move towards immune regulation to alter a fibrotic response towards wound healing, there is a clear need to account for biologic host variables such as age and sex to ensure the intended design is functional across populations. Although we did not investigate the underlying mechanisms of the immune difference (i.e. whether differences are due to hormone or chromosome complement), by incorporating age and sex as experimental variables, we show local immune responses unique to each fibrotic microenvironment and highlight the importance of biological factors in determining the fibrotic response. While effectively modulating specific immune populations may require interventions at different biological scales (e.g. gene, population, phenotype), our findings demonstrate that translatable biomaterials must be intentionally designed to account for host factors.

## Methods

### Ethics and Animal Study

All procedures were approved by the Johns Hopkins University (JHU) Animal Care and Use Committee. All mice were housed in JHU animal facilities under standard conditions with access to food and water ad libitum. Female and male C57BL6 mice aged 2 and 18 months were sourced from the NIA rodent colony under grant AG081564. The VML injury was performed bilaterally as previously described [42]. Mice were anesthetized (1.5-3% isoflurane), weighed, and received 3.25 mg/kg buprenorphine extended-release injectable suspension (Ethiqa-XR) subcutaneously for pain management. The hindlimb fur was then removed by shaving, and the surgical site was disinfected. A short, longitudinal incision was made in the skin and fascia overlaying the quadriceps femoris, then using sterile micro dissecting scissors and forceps a critical-sized defect (~3 mm x 3 mm x 3 mm) was excised from the muscle. The defect was then directly filled with sterile Dulbecco’s phosphate buffered saline (Saline, injury control), or polycaprolactone particulate (PCL, Polysciences, MW 50,000, 600 µm average size), or polyethylene particulate (PE, ultra-high MW, 125 µm average size).

Particulate was added using an 8 mm diameter curette (Moira 1121b) to place 25-30 mg of PCL or PE into the wound. The incision was then closed using sterile veterinary 5-0 nylon sutures and the wound plus material implant was repeated on the other hindlimb. Mice were placed in a clean cage with heat and closely monitored to ensure full recovery.

### Bulk Sequencing

#### Processing

A quadricep with material implant was collected immediately after euthanasia and stored in RNAlater at 4C for up to a week then −80C until further processing. Harvested tissues were then transferred to TRIzol (Invitrogen) and homogenized using a Bead Ruptor 12 (OMNI International) with ceramic beads (2.8 mm; OMNI International). RNA was isolated and purified using chloroform and RNeasy PLUS Mini Kit (Qiagen). Extracted RNA was submitted to Psomagen for QC, Library Prep, and bulk RNA-sequencing. Paired-end sequencing with a read depth of approximately 50 million reads per sample was performed using TruSeq stranded mRNA library Kit and run on an Illumina platform.

#### Quality Control and Alignment

Fastq level quality control was performed using fastp 0.20.1[63] and MultiQC v1.17[64]. Reads were then aligned using STAR 2.7.10a[65] with GRCh38 and GENCODE v29 annotations. The resulting reads were further inspected for outliers using principal component analysis (PCA) and correlation. Blinded variance stabilizing transformed (VST) values are used as inputs for both PCA and correlation. No outliers were identified or removed for subsequent analysis.

#### Differential Expression and Gene Set Enrichment Analysis

DESeq2 1.42.0[66] was used to perform differential expression using a negative binomial model with a Wald test to determine significance. Stratified pairwise comparisons were run, for example comparing male vs female separately for old only and young only samples and visa versa. Default DESeq2 metrics were used to filter genes based on expression levels and calculate normalization factors. All visualizations of gene expression use values derived from a blind variance stabilizing transformation of raw counts. Volcano plots of results were generated using EnhancedVolcano 1.26.0[67] with an adjusted P value threshold of 0.05 to indicate statistical significance and a foldchange threshold of +/-2. A select set of genes (*Mmp9, Pdpn, Cd80, Cd8a, Fn1, Themis3, Ly6g6e, Amd1, C7, Col4a4, Xist, Il1rl1, Ostn, Cd300e, Mmp19, Angptl1, Lyz1, Il10)* were labeled on volcano plots if significantly different to aid in visualization. Results were subsequently processed by gene set enrichment analysis using fgseaMultilevel from fgsea 1.20.0[68] with all Hallmark, KEGG, Matrisome gene sets using −log(p) * sign(fc) as the ranking metric where P is p-value and fc is fold-change. Multiple corrections for all gene sets were performed simultaneously.

### Flow Cytometry

#### Processing

Immediately after euthanasia, iLNs were separated from the inguinal fat pads and placed in 25mm HEPES in RPMI 1640 with L-Glutamine; once all iLN were collected they were then mashed through individual 70 µm cell strainer (Miltenyi Biotec) into a 15 mL conical with PBS. Quadriceps with material implant were collected after iLNs and the tissue was diced into 1-2 mm pieces then enzymatically digested under agitation for 45 minutes at 37C with 1.67 Wunsch U/mL (5mg/mL) Liberase TL (Roche Diagnostics), 0.2 mg/mL DNase I (Roche Diagnostics), 25 mM HEPES in RPMI 1640 medium with L-Glutamine (Gibco). After digestion samples were moved to ice and the digest quenched using cold 1% bovine serum albumin (BSA), 25 mM RPMI. For the pan immune panel, quadriceps were filtered through a 70 µm cell strainer followed by a 40 µm cell strainer, centrifuged, and rinsed twice in PBS before plating into a 96-well U-bottom plate for staining. For the ICS panel with stimulation, a Percoll density gradient was used. Briefly, cells were resuspended in an 80% Percoll solution, then 40% and 20% solutions were layered above before centrifuging at 2100 g for 30 minutes with acceleration and break set to 1. Cells were collected and washed twice with PBS before plating into a 96-well U-bottom plate. Plates were covered during staining to protect fluorophores.

#### Pan immune

Samples were centrifuged at 500g for 5 minutes and washed once with PBS prior to viability staining with Zombie NIR Fixable Viability for 30 mins on ice. Cells were then washed with staining buffer (1% BSA, 1mM EDTA in PBS) twice before surface marker staining for 45 minutes on ice. The control fluorescent minus one (FMO), single color controls (SCC), and master mix (MM) of the surface stain was made during the incubations steps. FMO, SCC, and MM were prepared using 1:20 dilution of TruStain FcX (BioLegend), 1:50 dilution of True-Stain Monocyte Blocker (BioLegend), and 1:50 dilution of Super Bright Complete Staining Buffer (eBioscience), antibodies in S.Table 1 were added at the stated concentrations and the staining buffer was used to bring to a final volume of 100 µL per well. For the MM, PE-Cy7 and APC-Cy7 were added to the MM immediately before adding the MM to the samples. After staining, cells were washed with buffer twice,then fixed in 100 µL of FluoroFix Buffer (Biolegend) for 15 minutes at room temperature. Plates were stored at 4C in staining buffer; immediately before data collection samples were washed in PBS and resuspended in a final 200 µL volume for samples and 100 µL for controls.

#### Intracellular Cytokine Staining (ICS)

Percolled samples were incubated in 1x eBioscience Cell Stimulation Cocktail Plus Protein Transport Inhibitors (eBioscience) diluted in Iscove’s Modified Dulbecco’s Media (IMDM, no phenol red) with L-Glutamine supplemented with 10% v/v heat-inactivated fetal bovine serum (Gibco) for 4 hours at 37C and 5% CO_2_. After stimulation, the plate was centrifuged and washed twice with PBS. Cells were stained with Zombie NIR for 30 minutes on ice, surface stained FMO, SCC, and MM were prepared using 1:20 dilution of TruStain FcX (BioLegend), 1:50 dilution of True-Stain Monocyte Blocker (BioLegend), and 1:50 dilution of Super Bright Complete Staining Buffer (eBioscience), antibodies in S.Table 2 were added at the stated concentrations and the staining buffer was used to bring to a final volume of 100 µL per well. After surface staining, samples were washed twice in PBS then fixed and permeabilized using Cyto-Fast Fix/Perm Solution (BioLegend) for 20 minutes at room temperature. Samples were then washed three times with 1x Cyto-Fast Perm Wash solution. For the cytokine FMO equivalent controls, the isotype for the antibody was used in its place. For all controls and MM, the FcX, Monocyte Blocker, Super Bright and antibodies at desired concentrations were resuspended in 1x Cyto-Fast Perm Wash solution and samples were stained for 20 minutes at room temperature. After staining, cells were washed once with Perm Wash then twice with 1% BSA in PBS. IL-17 is reported here as the sum of any CD4+ cell expressing either IL-17a or IL-17f.

#### Flow Cytometry and Data Analysis

Prior to data collection samples were washed and resuspended in PBS and with a final 200 µL volume for samples and 100 µL for controls. All flow cytometry data were collected on the 4-laser Cytek Aurora (V, B, YG, R) with automated sample loader spectral cytometer. Spectral unmixing was performed using SpectroFlo (Cytek Biosciences v3.3), autofluorescence signatures were extracted from unstained samples and included as independent fluorophores. Manual flow gating to identify cell populations of interest was performed using FlowJo software (Tree Star, v10.9), UMAP dimensional reductions were completed using the FlowJo plugin[69]. Counts are calculated per quadricep by taking the instrument counts for a known collected volume and calculating total cells per 200 µL.

### Histology

#### Sample Preparation

Muscles were excised and fixed for 48 hours in 10% neutral buffered formalin with agitation at room temperature. Tissues were then dehydrated using an ethanol series (70%, 80%, 95%, 100%) before clearing for 1.5 hours in xylenes. Samples were then stored in melted paraffin overnight at ~60C. Tissues were embedded in paraffin blocks and sectioned at 7 µm thickness through the Johns Hopkins Oncology Tissue Services SKCCC core facility. Masson’s trichrome staining was completed by the core facility. Picrosirius Red staining was completed following manufacturer protocols (Abcam ab150681).

#### Immunofluorescence

Slides were deparaffinized in xylenes and rehydrated in reverse ethanol concentrations (100%, 95%, 80%, 70%, followed by Type 1 water). AR-6 (Akoya Biosciences) was used for heat-induced epitope retrieval in a steamer for 15 minutes. Exogenous peroxidase was quenched using 3% H_2_O_2_ (Sigma) for 15 minutes. Slides were then blocked in 10% BSA, 0.05% Tween 20 in PBS for 30 minutes. Primary antibodies, S.Table 3, were incubated for 30 minutes at room temperature, slides were then rinsed in 1x TBST three times, incubated for 30 minutes rabbit-on-mouse IgG horse radish peroxidase polymer (Biocare Medical), rinsed in 1x TBST 3x then reacted with tyramide signal amplification reagents and Opals for 10 minutes. Slides were then stripped by heat (steamer) with AR6 for 15 minutes before repeating the staining process for the next primary antibody. Nuclei were counterstained with 4’,6-diami-dino-2-phenylindole (Spectral DAPI, Akoya) for 5 min before mounting using DAKO mounting medium (Agilent). Imaging was performed on an Axio Imager.A2 (Carl Zeiss Microscopy, LLC) with Zen software (Zeiss). Images were linearly contrasted with isotype or primary delete controls for each channel setting the contrast levels. Images were quantified using QuPath-0.5.1. To quantify within the FBR, an ROI was drawn around fibrosis only, a mask was created on the DAPI channel using a pixel classifier (settings were adjusted to outline just the tissue without the empty particle areas), cells were identified with Cell Detection, positive cell detection was quantified using an object classifier and vessels sizes were quantified using a pixel classifier (minimum area was set to 60 um^2^).

#### Masson’s Trichrome Analysis

QuPath-0.5.1 was used to quantify the collagen (blue regions) in Masson’s trichrome using a trained pixel classifier (Fig. S3B). Briefly 3-5 ROIs of the collagen, cells, and particulate location were manually annotated as collagen, tissue, or ignore (empty space). For our images the artificial neural network (ANN_MLP) classifier with full resolution and default multiscale features worked best to identify the blue collagens versus the tissue regions. An ROI was drawn within the material fibrosis only, excluding the muscle and any fat attached above the particulate, and then the pixel classifier was run within that ROI only. The data is presented as percent collagen for the area within the fibrosis.

#### Picrosirius Red (PSR) Analysis

QuPath-0.5.1 was used to analyze PSR as the percent red or ratio red to green within the fibrosis, analysis pipeline is described visually (Fig. S4B–C). An ROI was drawn around the particulate induced fibrosis only, ignoring the muscle and any fat that may still be attached to the tissue. A pixel threshold (moderate, gaussian, gamma = 1) was set on a brightfield channel to identify the fibrosis ROI around the empty particulate. Two full resolution pixel thresholds (gaussian, gamma = 0) were set to identify both red and green fiber pixels within each respective channel. For the PE particulate, since the PE is still present on the slides and the particulate appears in all 3 channels (red, green, blue), the blue channel was used to subtract the particulate area from the fibrosis ROI before proceeding with the red and green analysis. The red/green is presented as percentage of the area identified as fibrosis.

### Statistics

Except for bulk RNA sequencing, all statistical analyses were performed utilizing Graphpad Prism v10 and a p-value of less than 0.05 was considered statistically significant unless otherwise stated. All data within a plot were collected simultaneously and analyzed using a two-way ANOVA with Sidak posthoc analysis for the four main comparison (young female to young male; young to aged female; young to aged male; aged male and aged female).

## Acknowledgements

We would like to thank the JHU Oncology Tissue Services SKCCC core facility for histology, sectioning, and performing Masson’s trichrome staining. This study was supported by the National Institutes of Health under award numbers R01AG082965, DP1AR076959, and R01EB028796, NSF Graduate Research Fellowship Program DGE1746891(AR) and K99AG081564 (JCM).

## Data and code availability

Raw FASTQ files and aligned counts files will be made available for download through the National Center for Biotechnology Information (NCBI) Gene Expression Omnibus (GEO). GEO accession numbers will be provided once available.

## Author contributions

JCM, AR, ASR, KS, SK, KK, NR, LDH performed experiments and interpreted results. JCM, AR, KS, FH developed and performed flow cytometry. CC and KK performed computational analysis. All authors have read and approved the manuscript

