## Supplemental Data for "Age and sex alter the immune response in a chronic fibrosis model via changes in T cell and macrophage phenotype"

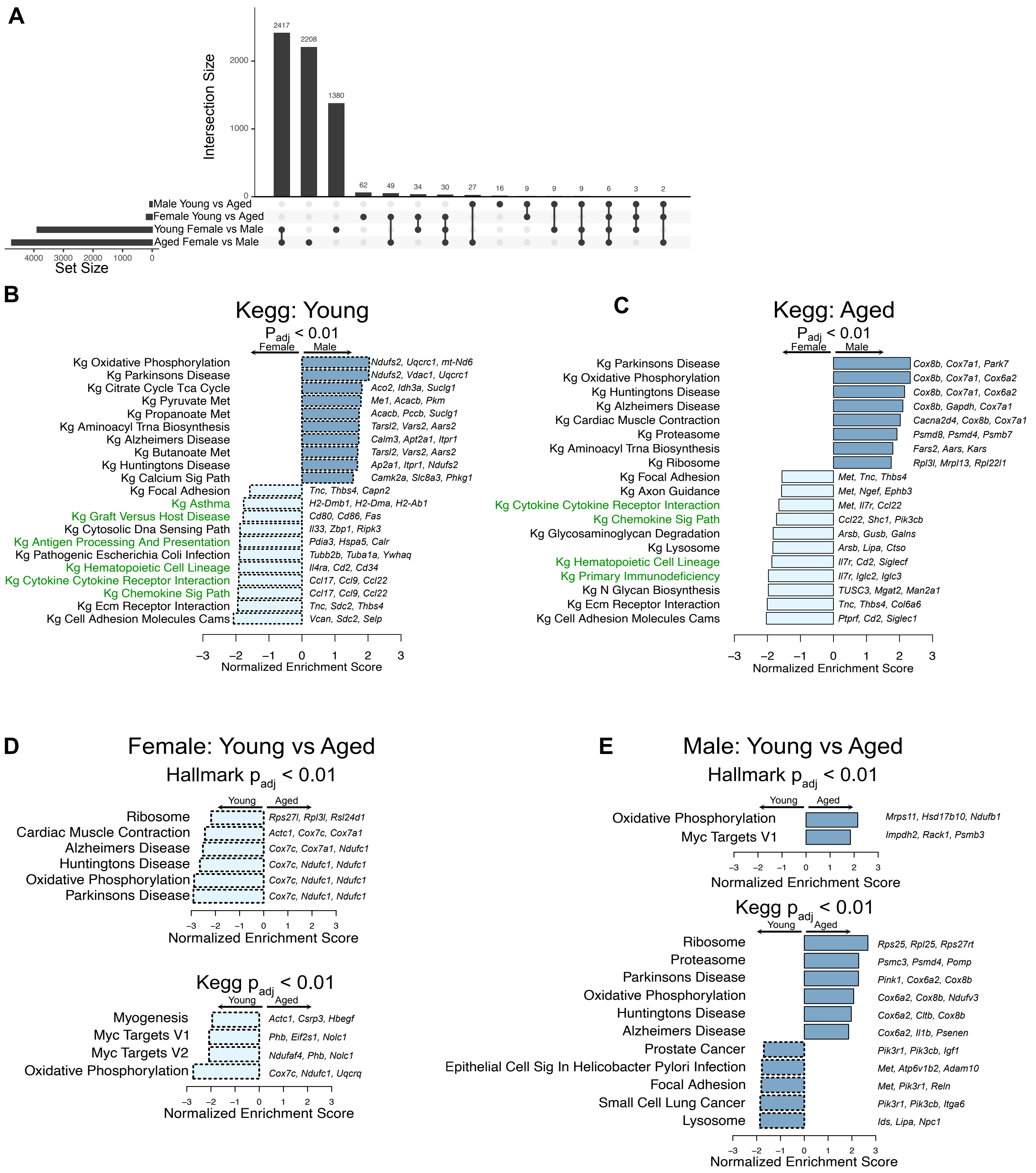


Supplemental Figure 1: (A) Overlap and independence of DEG sets for each of the 4 comparisons visualized by upset plot. (B-C) Kegg GSEA pathways with an adjusted p value < 0.01 for comparisons by sex. (D-E) Hallmark and Kegg GSEA for the comparisons by age. Dashed lines indicate GSEA pathways up in young and solid lines indicate up in aged.

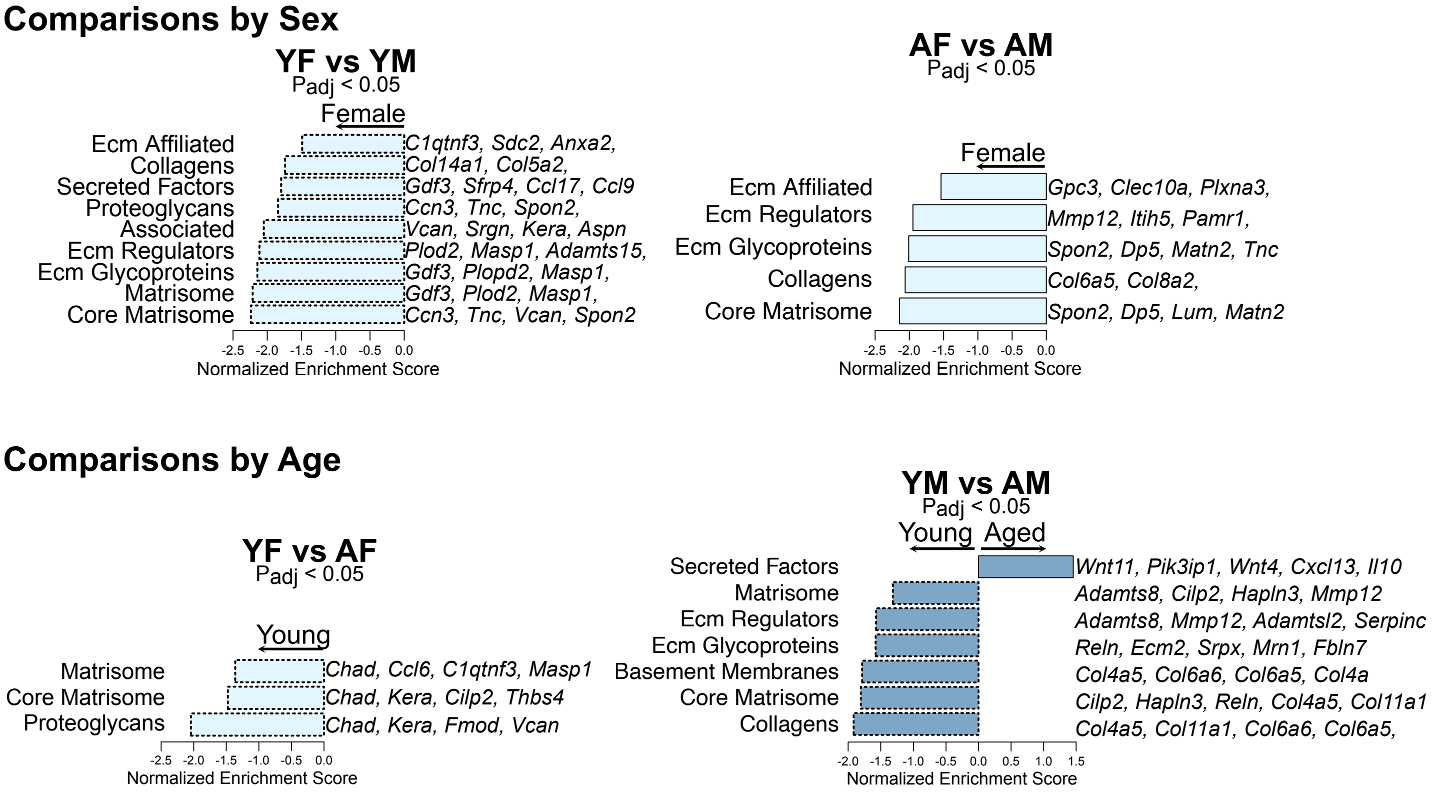


Supplemental Figure 2: Naba Matrisome GSEA with Padj < 0.05 for each of the four comparisons.

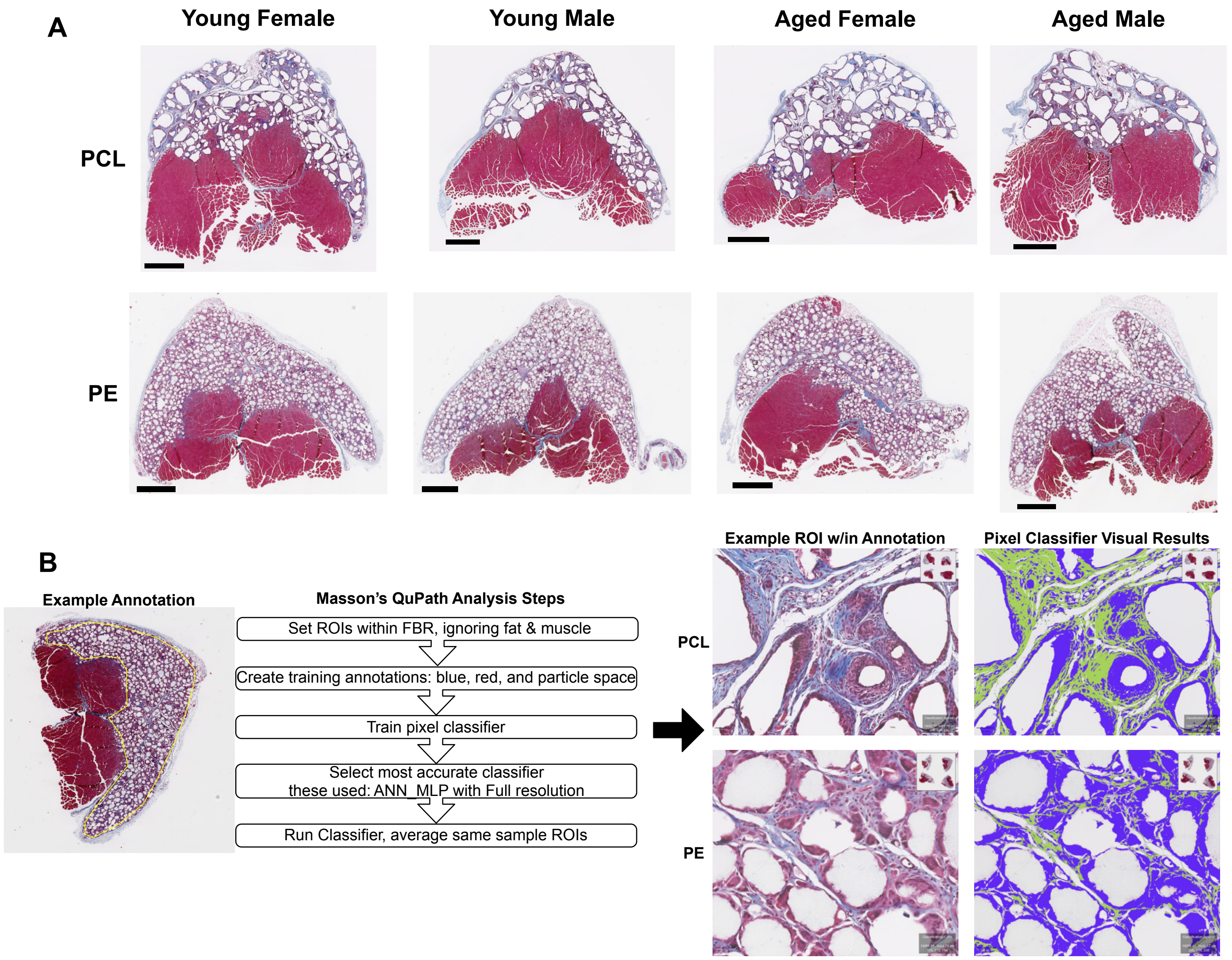


Supplemental Figure 3: (A) Representative Masson's Trichrome staining for each sample type, scale bar = 1 mm. (B) QuPath analysis steps for Masson's Trichrome staining to be able to quantify the percentage of the FBR that stained blue and representative images of the results showing the classifier identifying the collagens (yellow-green) verses the rest of tissue (dark blue).

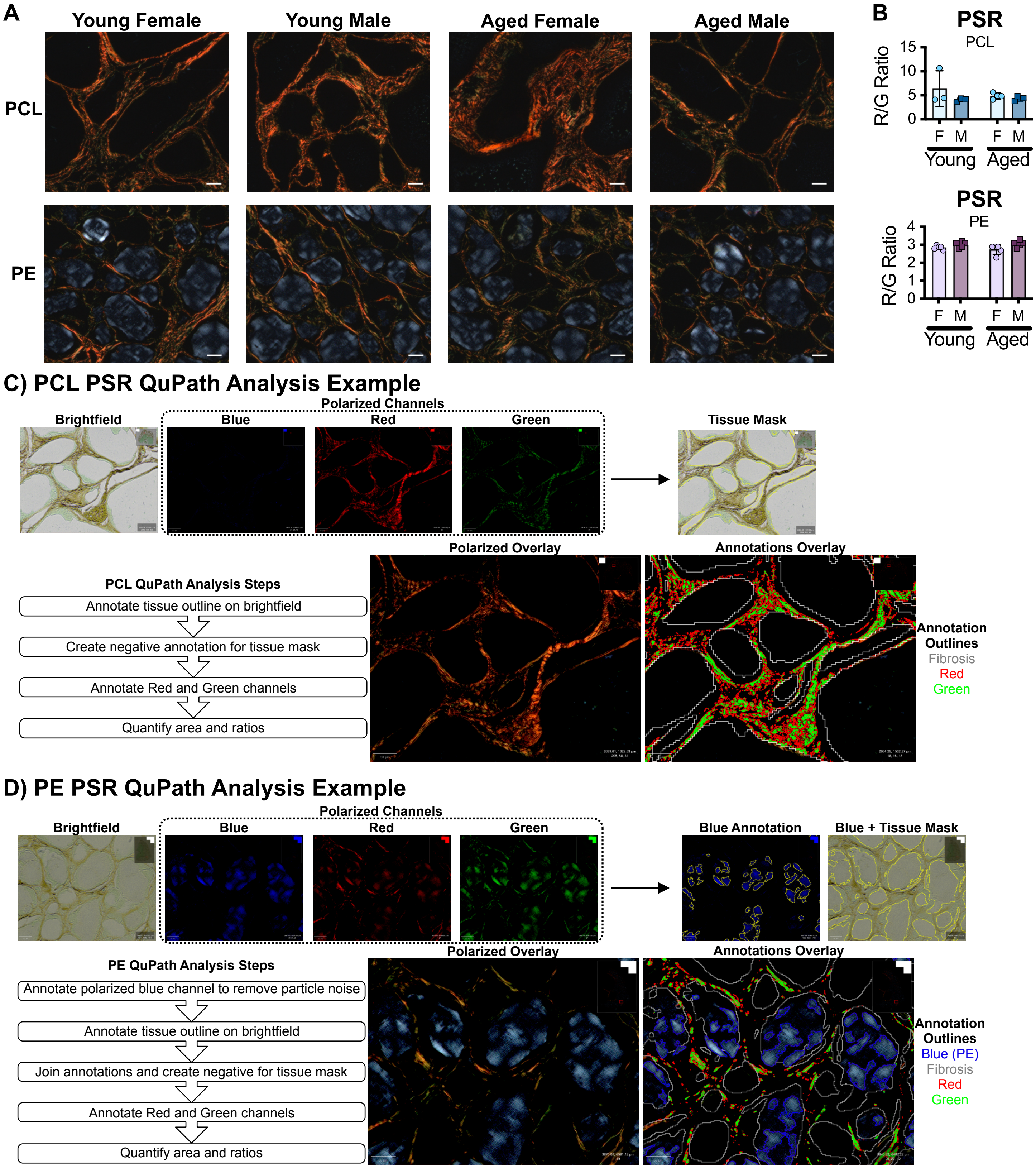


Supplemental Figure 4: (A) Red:green ratio for picrosirius (PSR) quantification. (B) Representative polarized images for each sample (C-D) QuPath analysis steps and images to show how annotations were used to mask the tissue area and then the red and green pixel annotations of the polarized channels. PCL differed from PE in the blue channel was used to subtract the PE particles that remained on the samples and appeared in all 3 polarizer channels.

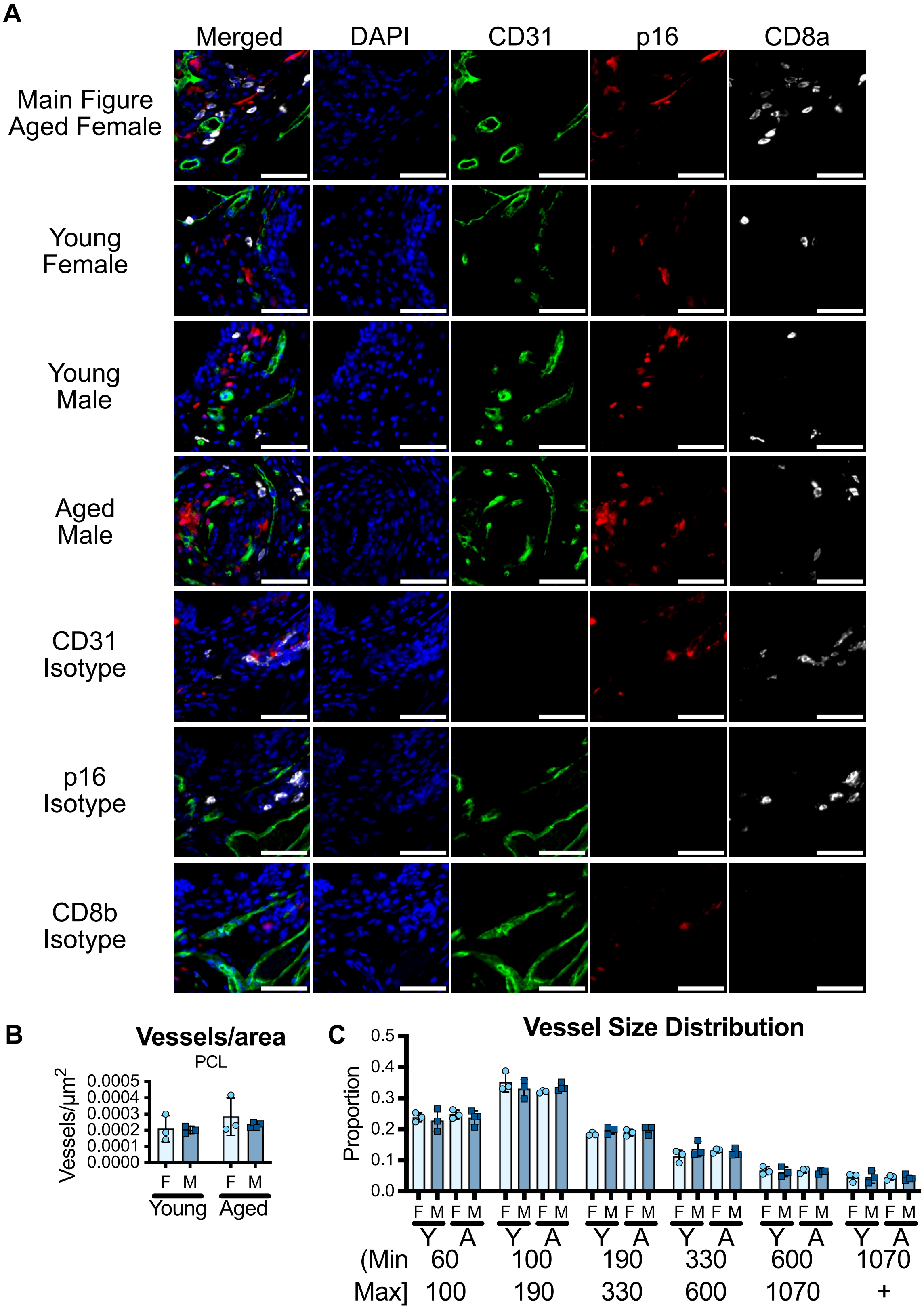


Supplemental Figure 5: (A) Representative images of all age and sex groups with corresponding single channel images and isotype controls for Figure 2E. Scale bars = 50 µm. All images acquired on a Zeiss Axio Imager A2 and linearly contrasted identically in Zen. (B) QuPath pixel classification of the number vessels within the FBR area (C) Proportion of vessels distributed by vessel size. Statistics: 2 way ANOVA with Sidak posthoc, adjusted p = *0.05, **0.01, ***0.001, ****<0.0001.

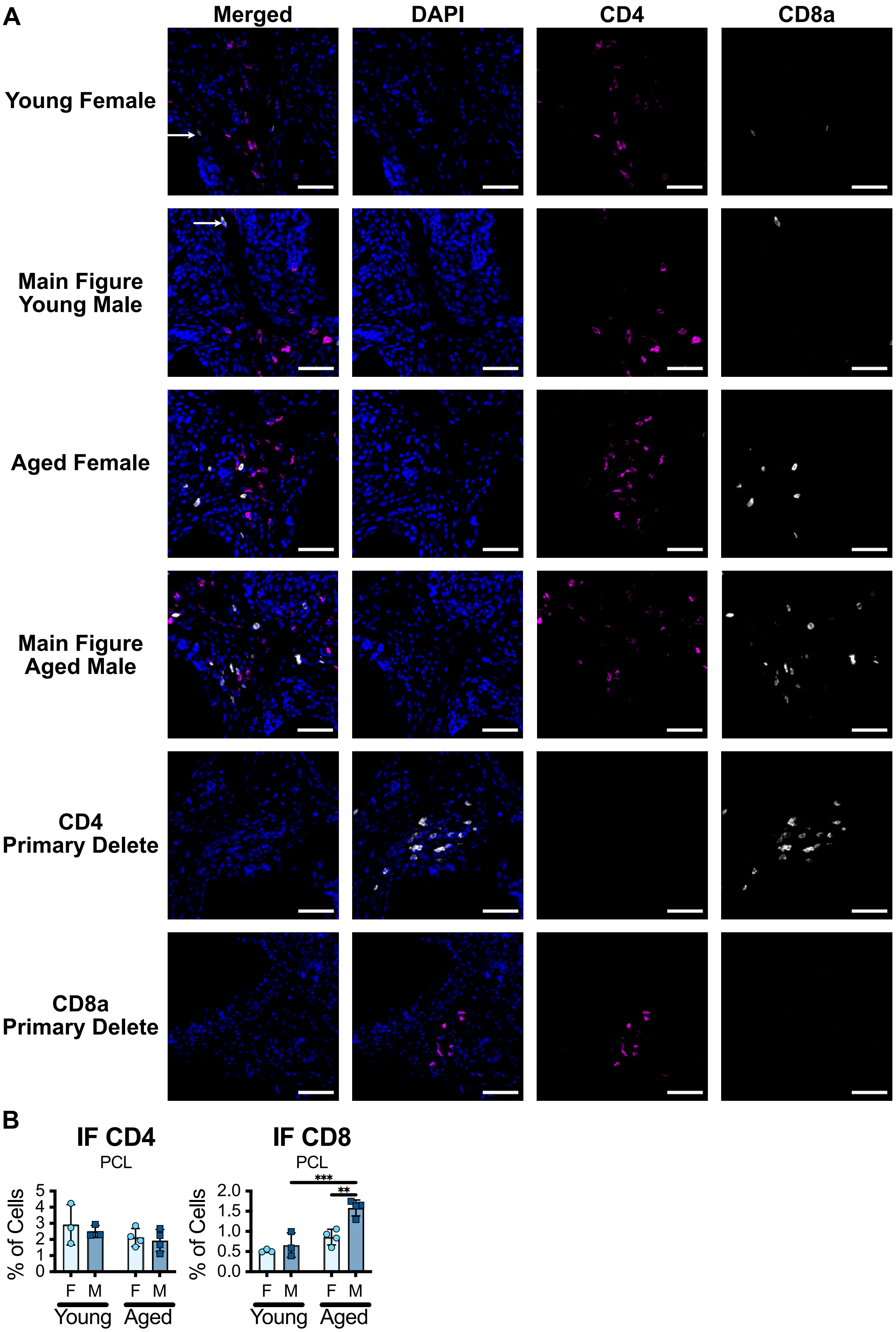


Supplemental Figure 6: (A) Representative images for both age and sexes with corresponding single channel images and isotype controls for Figure 4A. Scale bars = 50 μm. All images acquired on a Zeiss Axio Imager A2 and linearly contrasted identically.  White arrow on young point to CD8 within the ROI.(B) Quantification of CD4 and CD8 cells within the FBR. Statistics: 2 way ANOVA with Sidak posthoc analysis comparing only YF-YM, YF-AF, YM-AM, AF-AM, adjusted p = *0.05, **0.01, ***0.001, ****<0.0001.

Supplemental Table 1: Pan Immune Flow Cytometry Panel

| Fluorophore | Marker | Clone | Catalog | Manufacturer | RRID | Dilution |
|---|---|---|---|---|---|---|
| BV421 | CD86 | GL-1 | 105032 | BioLegend | AB_2650895 | 200 |
| Super Bright 436 | CD19 | 1D3 | 62-0193-82 | ThermoFisher | AB_2688107 | 100 |
| Pacific Blue | Ly6G | 1A8 | 127612 | BioLegend | AB_2251161 | 250 |
| BV480 | SiglecF | E50-2440 | 746668 | BD Optibuild | AB_2743940 | 100 |
| BV605 | CD45 | 30-F11 | 103140 | BioLegend | AB_2562342 | 300 |
| BV650 | Ly6C | HK1.4 | 128049 | BioLegend | AB_2800630 | 1200 |
| BV711 | GD TCR | GL3 | 563994 | BD | AB_2738531 | 200 |
| BV750 | B220 | RA3-6B2 | 103261 | BioLegend | AB_2734157 | 200 |
| BV785 | F480 | BM8 | 123141 | BioLegend | AB_2563667 | 300 |
| BB515 | cKIT | 2B8 | 564481 | BD | AB_2738826 | 500 |
| Spark Blue 550 | CD3 | 17A2 | 100260 | BioLegend | AB_2832258 | 100 |
| PerCP | MHCII | M5/114.15.2 | 107624 | BioLegend | AB_2191073 | 200 |
| BB700 | CD8a | 53-6.7 | 566409 | BD | AB_2744467 | 100 |
| PE | CD301b | MGL2 | 146804 | BioLegend | AB_2562944 | 500 |
| PE-Dazzle 594 | CD11c | N418 | 117348 | BioLegend | AB_2563655 | 500 |
| PE-Cy7 | CD200R3 | Ba13 | 142212 | BioLegend | AB_2814046 | 400 |
| APC | CD206 | C068C2 | 141708 | BioLegend | AB_10900231 | 200 |
| AF647 | NK1.1 | PK136 | 108720 | BioLegend | AB_2132713 | 200 |
| AF700 | CD11b | M1/70 | 101222 | BioLegend | AB_493705 | 400 |
| Zombie NIR | Viability |  | 423106 | BioLegend |  | 3000 |
| APC-Fire 750 | CD9 | MZ3 | 124814 | BioLegend | AB_2783073 | 500 |
| APC-Fire 810 | CD4 | GK1.5 | 100480 | BioLegend | AB_2860583 | 100 |
| FluoroFix 100 µL for 15 min @ RT |  |  | 422101 | BioLegend |  |  |

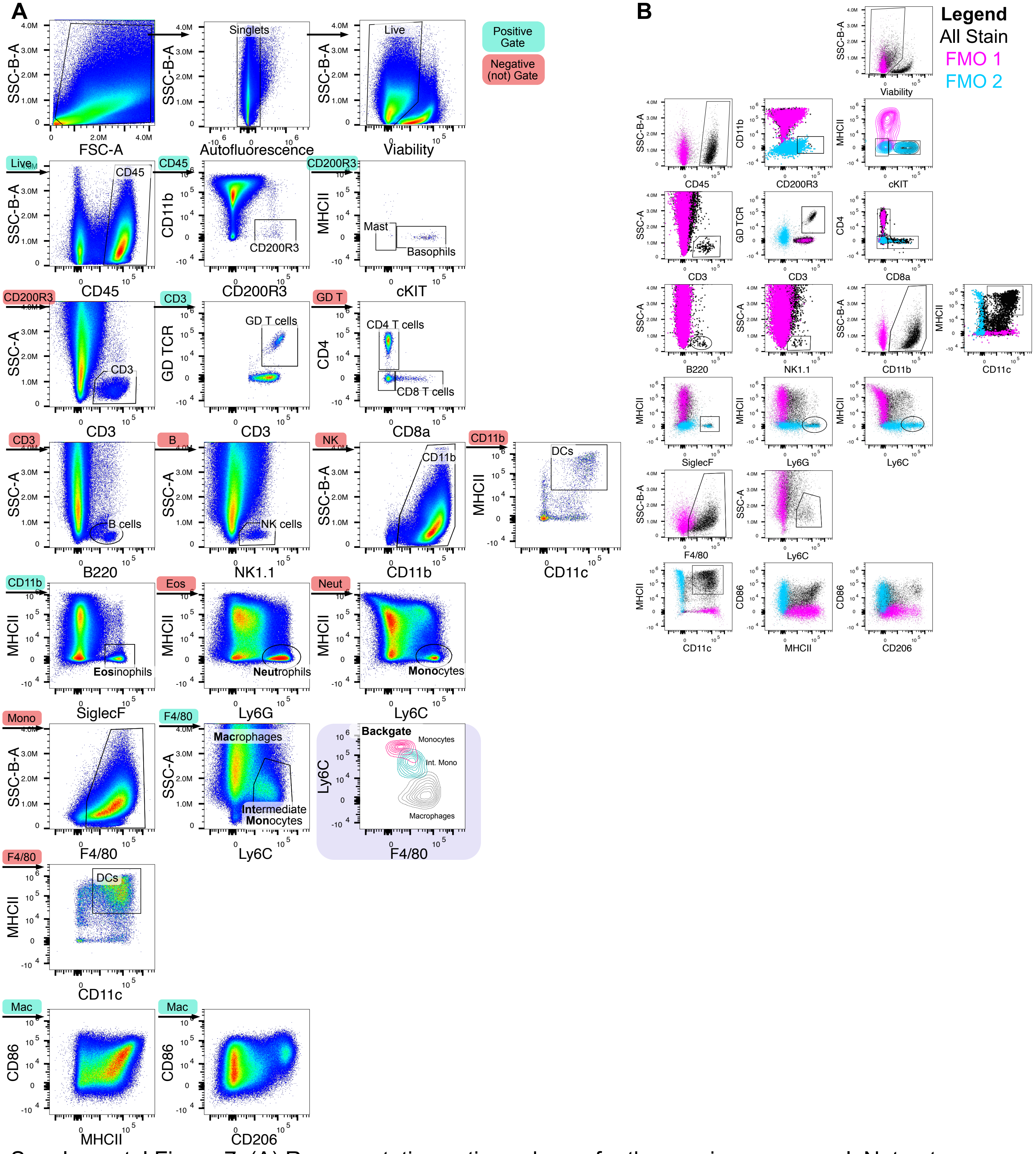


Supplemental Figure 7: (A) Representative gating scheme for the pan-immune panel. Not gates were used to remove cell populations from downstream gates and are identified by a red background above the arrow. Standard positive gates are identified in green. The monocyte, intermediate monocytes, and macrophage populations were backgated (purple outline) to show the Ly6C to F4/80 shift of those myeloid populations. (B) FMO control overlay of the gating scheme.

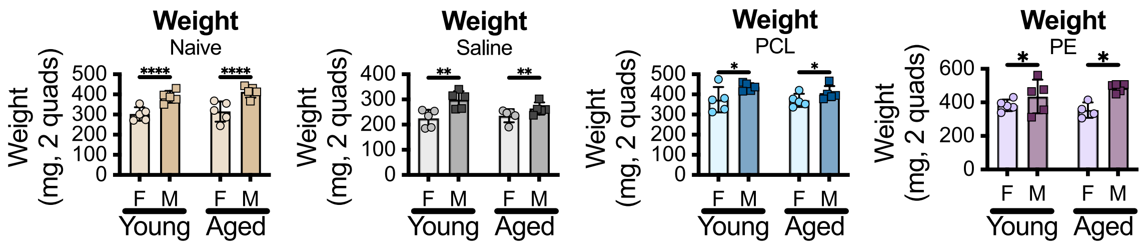


Supplemental Figure 8. Weights of two quadriceps for naïve and 6-week VML- saline, -PCL, and -PE tissue samples. Statistics: 2 way ANOVA with Sidak posthoc analysis comparing only YF-YM, YF-AF, YM-AM, AF-AM, adjusted p = *0.05, **0.01, ***0.001, ****<0.0001.

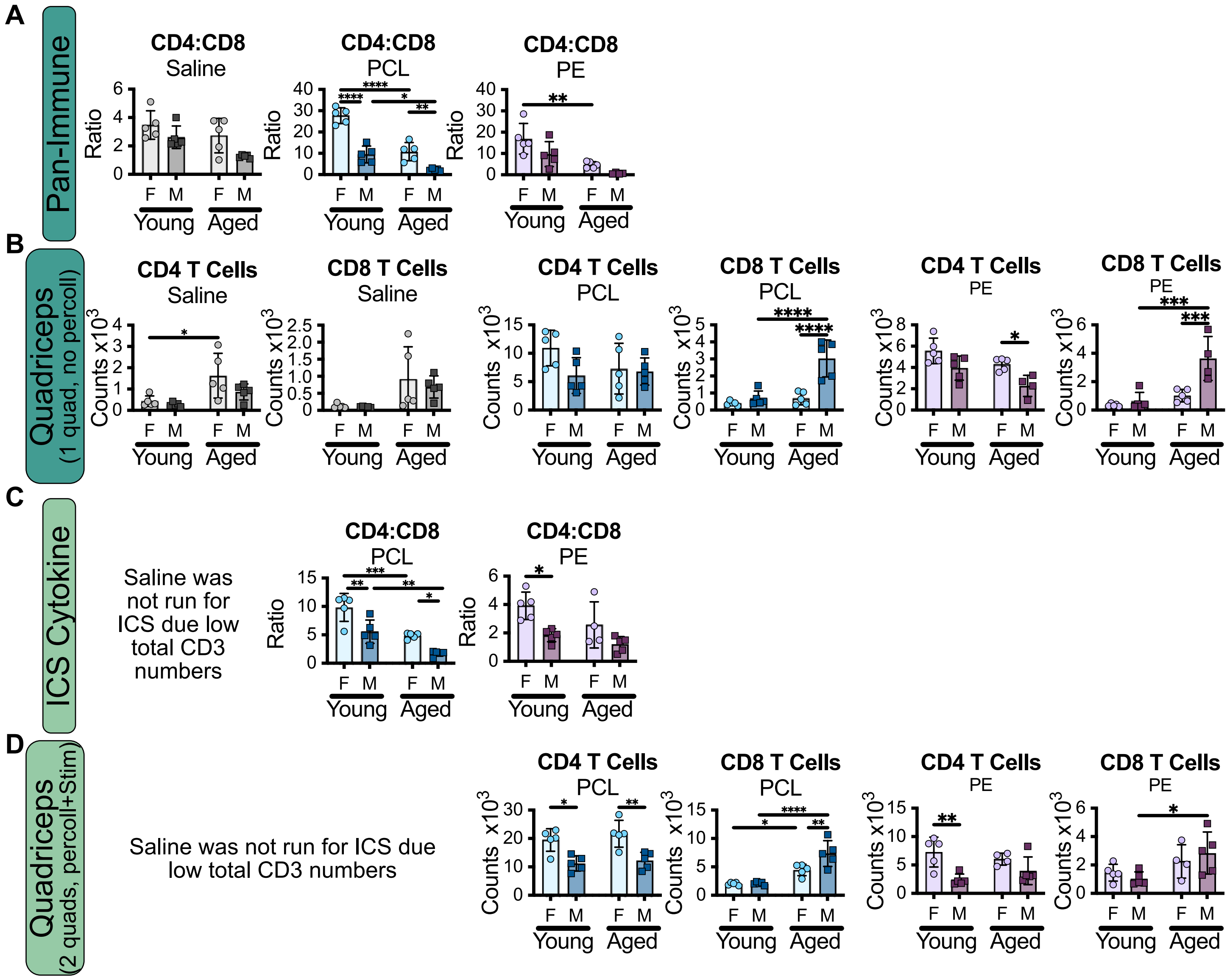


Supplemental Figure 9: (A) CD4:CD8 T cell ratios from the two different cell processing panels the pan-immune is stained on a whole quadricep after digest while the ICS Cytokine is from two quadriceps that are digested, percolled, and stimulated for 4 hours. (B) CD4 and CD8 T cell counts from quadriceps processed for the pan-immune and ICS Cytokine. Statistics: 2 way ANOVA with Sidak posthoc analysis comparing only YF-YM, YF-AF, YM-AM, AF-AM, adjusted p = *0.05, **0.01, ***0.001, ****<0.0001.

Supplemental Table 2: Intracellular Staining Cytokine Panel

| Fluorophore | Marker | Clone | Catalog | Manufacturer | RRID | Dilution | Step |
|---|---|---|---|---|---|---|---|
| BV421 | TCRb | H57-597 | 109230 | BioLegend | AB_2562562 | 200 | Surface |
| BV510 | CD45 | 30-F11 | 103138 | BioLegend | AB_2563061 | 200 | Surface |
| BV605 | CD4 | GK1.5 | 100451 | BioLegend | AB_2564591 | 150 | Surface |
| BV650 | CD11b | M1/70 | 101259 | BioLegend | AB_2566568 | 400 | Surface |
| BV711 | GD TCR | GL3 | 563994 | BD | AB_2738531 | 200 | Surface |
| BV786 | IL4 | 11b11 | 564006 | BD | AB_2738538 | 50 | Intracellular |
| Vio B515 | IL10 | REA1008 | 130-116-968 | Miltenyi | Not Available | 100 | Intracellular |
| Spark Blue 550 | CD3 | 17A2 | 100260 | BioLegend | AB_2832258 | 100 | Surface |
| BB700 | CD8a | 53-6.7 | 566409 | BD | AB_2744467 | 100 | Surface |
| PE | IL17f | 9D3.1C8 | 517008 | BioLegend | AB_10690818 | 200 | Intracellular |
| PE-eFluor 610 | IL13 | eBio13A | 61-7133-82 | ThermoFisher | AB_2574654 | 200 | Intracellular |
| PE-Cy5 | IL2 | JES6-5H4 | 503824 | BioLegend | AB_2123674 | 500 | Intracellular |
| APC | IFNg | XMG1.2 | 505810 | BioLegend | AB_315404 | 150 | Intracellular |
| AF647 | NK1.1 | PK136 | 108720 | BioLegend | AB_2132713 | 200 | Surface |
| AF700 | IL17a | TC11-18H10.2 | 506914 | BioLegend | AB_536016 | 200 | Intracellular |
| Zombie NIR | Viability | | 423106 | BioLegend | | 5000 | Surface |
| BV786 | R IgG1 | R3-34 | 563847 | BD Biosciences | AB_2869525 | Concentration match to cytokine | Isotype |
| Vio B515 | H IgG1 | REA293 | 130-114-556 | Miltenyi | Not Available | | Isotype |
| PE | M IgG1 | MOPC-21 | 400112 | BioLegend | AB_2847829 | | Isotype |
| PE-eFluor 610 | R IgG1 | eBRG1 | 61-4301-82 | ThermoFisher | AB_2637348 | | Isotype |
| PE-Cy5 | R IgG2b | RTK4530 | 400609 | BioLegend | AB_326553 | | Isotype |
| APC | R IgG1 | RTK2071 | 400412 | BioLegend | AB_326518 | | Isotype |
| AF700 | R IgG1 | RTK2071 | 400420 | BioLegend | AB_493781 | | |
| Cyto-Fast Fix/Perm Buffer Set | | | 426803 | BioLegend | | | |
| Isotype key: M = mouse, R = rat, H = human | | | | | | | |

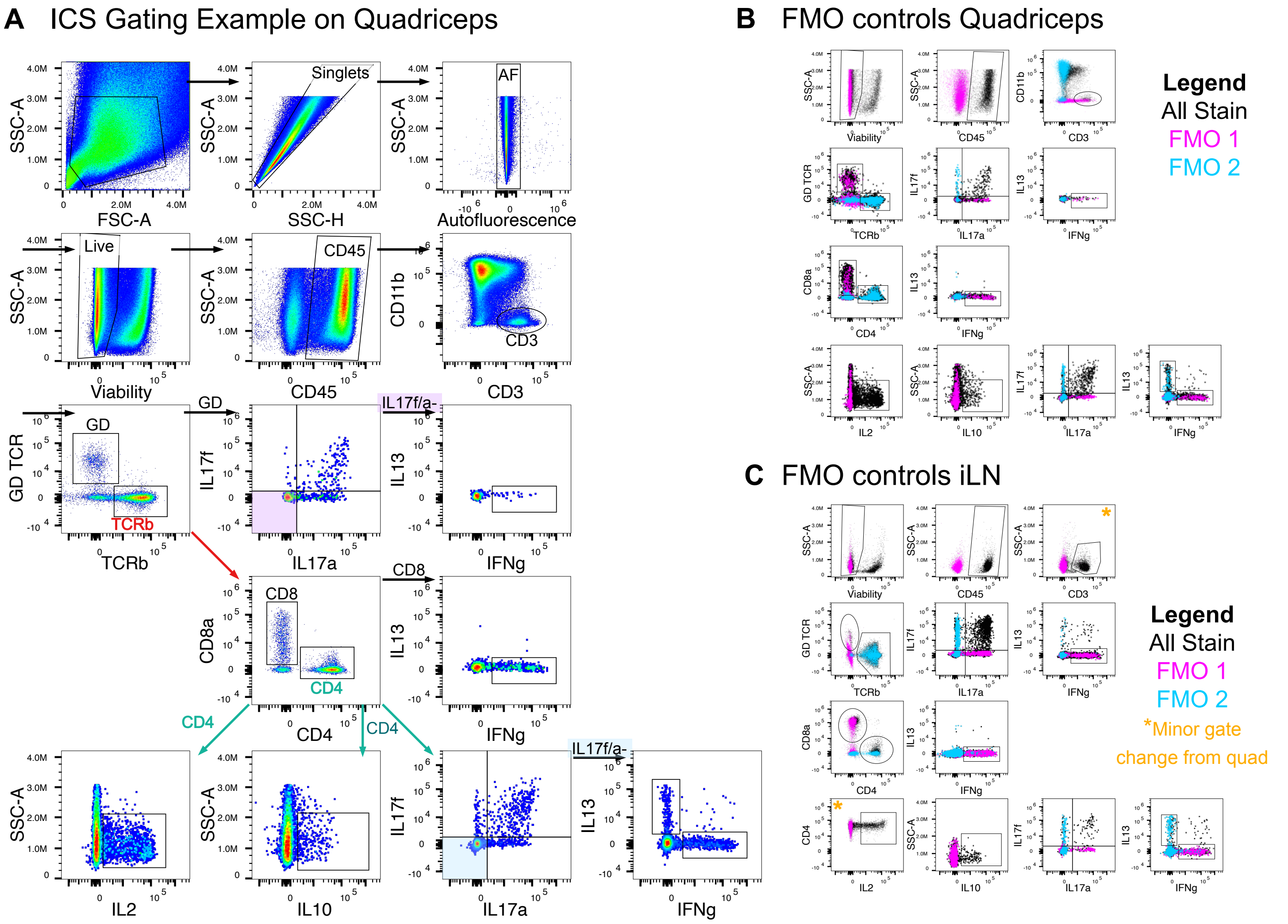


Supplemental Figure 10: (A) Representative gating scheme for the intracellular staining (ICS) panel. Populations are identified as: GD T cells (CD3+GD TCR+ TCRb-), CD8 T cells (CD3+GD-TCRb+ CD8a+CD4-), CD4 T cells (CD3+GD-TCRb+CD8a-CD4+). For GD cytokines IL17a and IL17f were first gated then from the double negative the proportion of IFNg were gated. For CD8 T cells only IFNg was gated. For CD4 T cell Cytokines as IL2 and IL10 are not specific to T helper phenotype they were gated from the parent CD4 population. For the CD4 T helper cytokines: IL17a and IL17f were gated on all CD4s then IFNg and IL13 were gated from the double negative IL17a/f. (B) Fluorescence minus one staining controls (FMOs) were used to identify cytokine populations. For each of the cytokines the respective isotype controls were used in place of the antibody (ie the IFNg FMO included APC Rat IgG1). (C) FMOs and gating schematic for inguinal lymph nodes (iLN); gating followed the layout of the quadriceps with minor variations in the CD3 gate vs SSC and IL2 vs CD4.

A

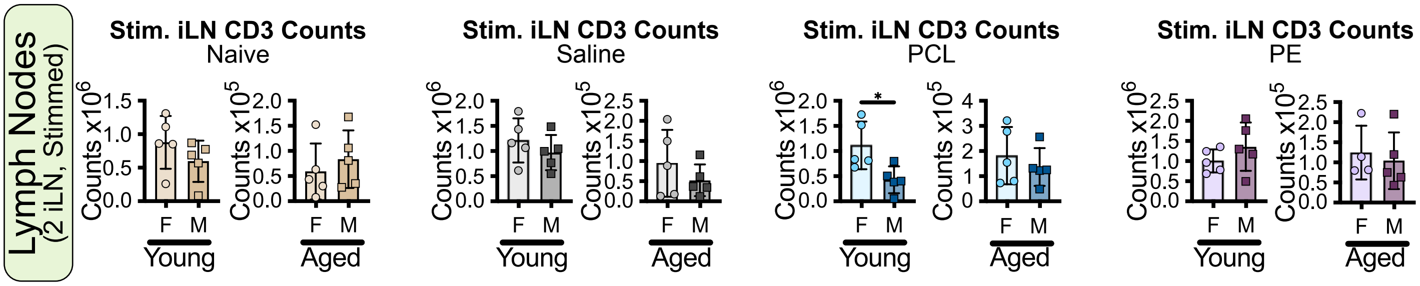


B

| | PCL x10³ | | PE x10³ | |
|---|---|---|---|---|
| | **Pan Immune** | **ICS** | **Pan Immune** | **ICS** |
| **Young Female** | 15.5 ± 4.4 | 15.9 ± 3.3 | 7.9 ± 1.7 | 7 ± 2.3 |
| **Young Male** | 10.0 ± 5.0 | 10.5 ± 2.1 | 5.8 ± 1.5 | 3.2 ± 1.3 |
| **Aged Female** | 12.2 ± 6.7 | 19.6 ± 4.3 | 7.5 ± 1.7 | 7.5 ± 1.5 |
| **Aged Male** | 15.3 ± 4.8 | 14.4 ± 2.4 | 8.6 ± 2.2 | 6.2 ± 2.0 |
| Calculated: Mean count +/- SD for one quad | | | | |

Supplemental Figure 11: (A) CD3 T cell calculated cell counts for Naïve, 6-week VML- saline, -PCL, from two inguinal lymph nodes after 4 hours of stimulation. Young and aged are plotted independently as there is a fold difference in values. Outlined plots are shown on the same axis within the main text. (B) Table comparison of calculated number of T cells between the two digest and staining methods. Statistics: 2 way ANOVA with Sidak posthoc analysis comparing only YF-YM, YF-AF, YM-AM, AF-AM, adjusted p = *0.05, **0.01, ***0.001, ****<0.0001.

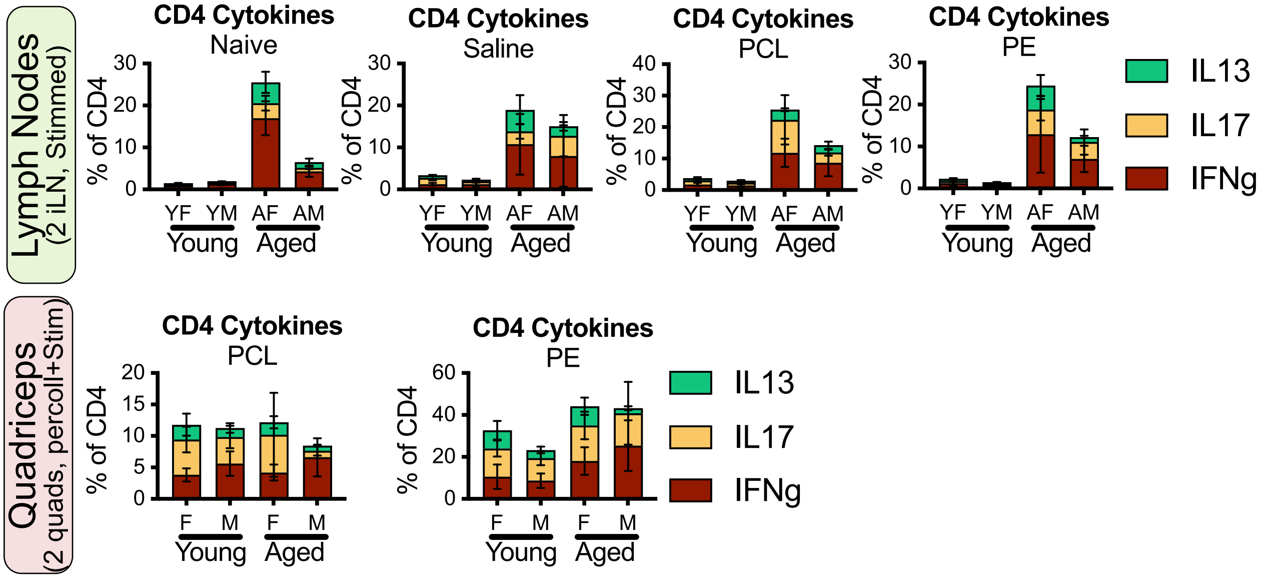


Supplemental Figure 12: (A) Overlay of proportions of IL13 (green), IL17 (yellow) and IFNg (red) proportions in CD4 T cells of the inguinal lymph nodes (no statistics shown).

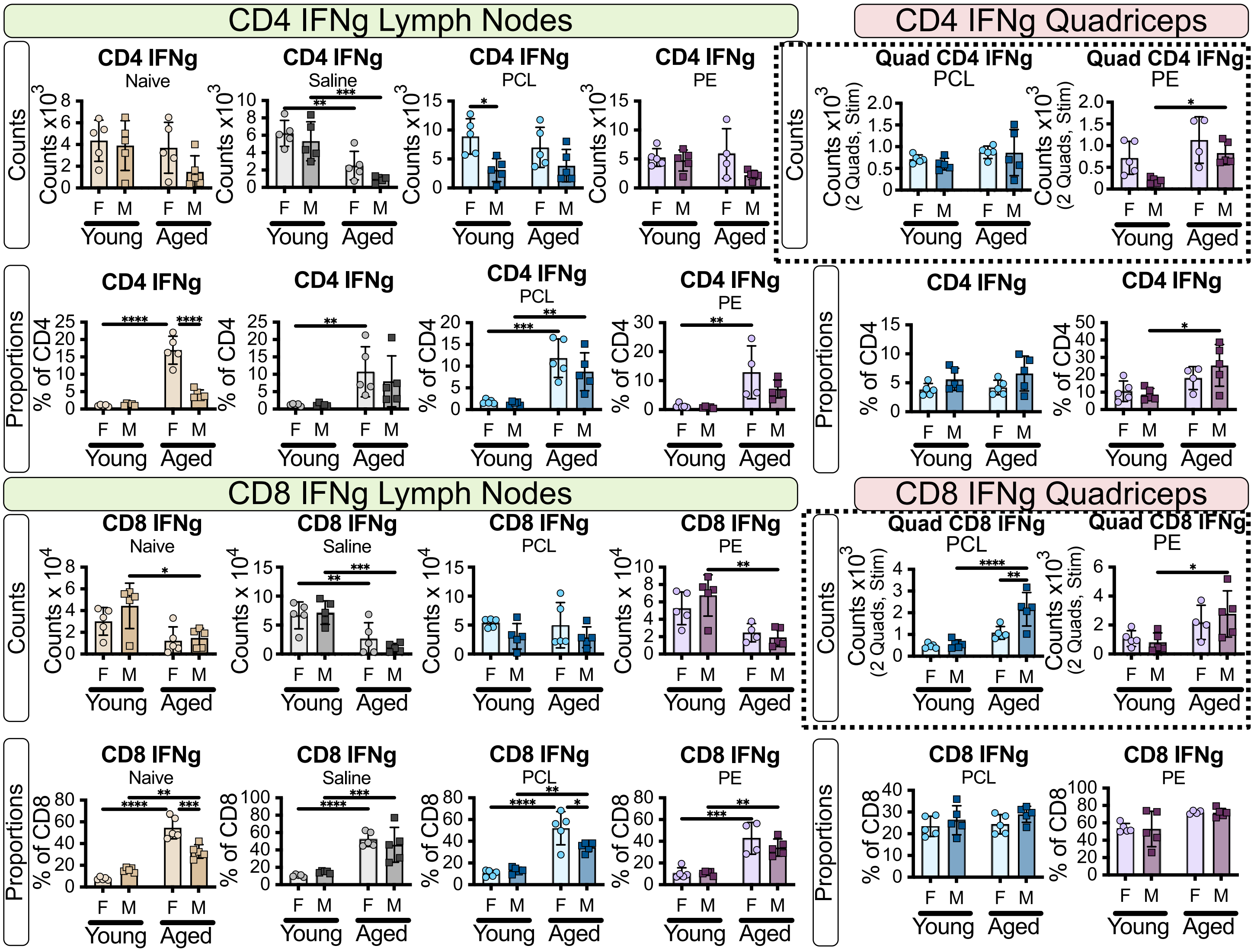


Supplemental Figure 13: Counts and proportions of CD4 and CD8 IFNg. Dashed outline indicates plots used in main figure, repeated here for easy comparison. Statistics: 2 way ANOVA with Sidak posthoc analysis comparing only YF-YM, YF-AF, YM-AM, AF-AM, adjusted p = *0.05, **0.01, ***0.001, ****<0.0001.

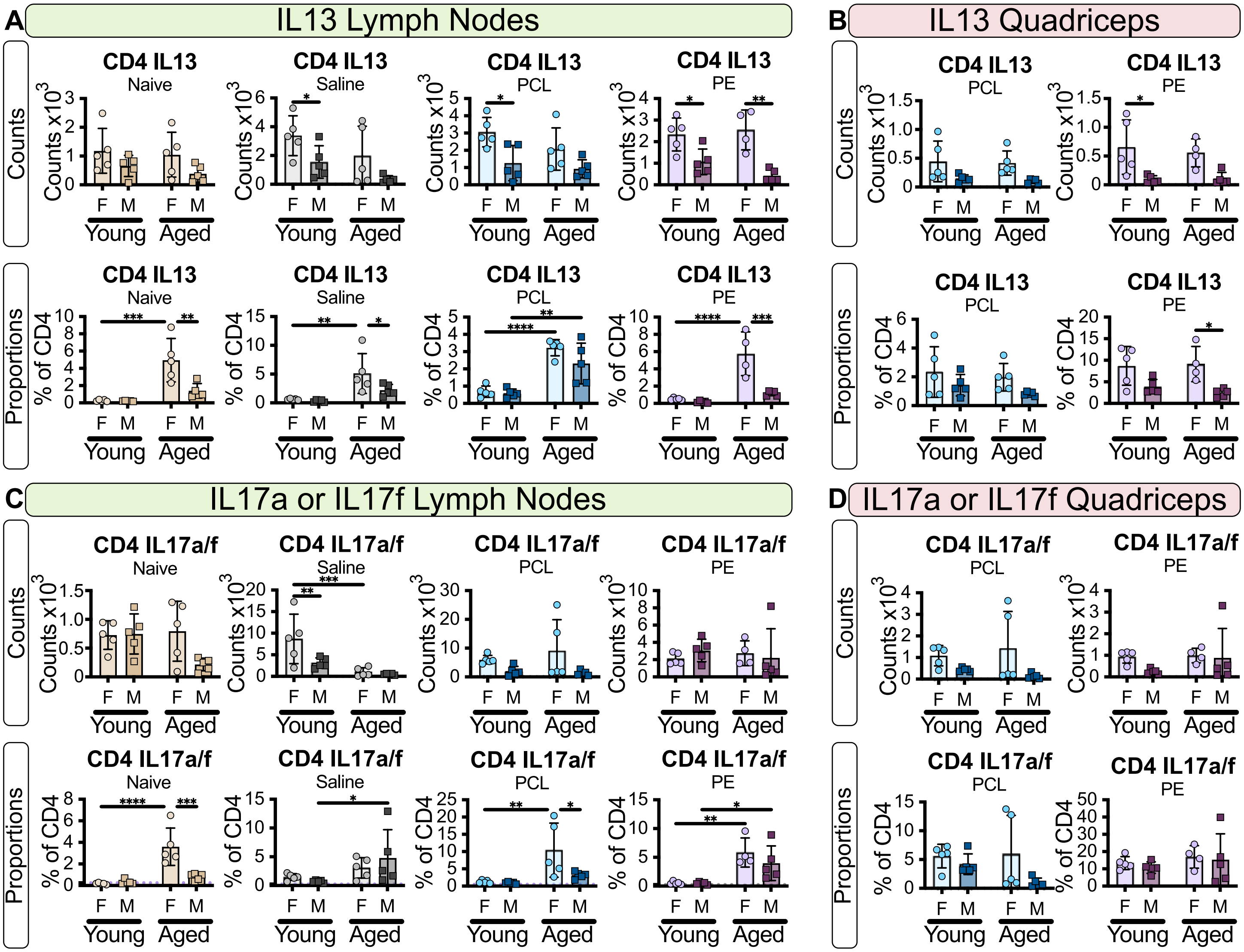


Supplemental Figure 14: (A) Overlay of proportions of IL13 (green), IL17 (yellow) and IFNg (red) proportions in CD4 T cells of the inguinal lymph nodes (no statistics shown). (B) Proportion and counts of IL13 expression in CD4 T cells from iLN and Quadriceps (C) Proportions and counts of any combination of IL17a and/or IL17f expression in CD4 T cells from iLN and quadriceps. Statistics: 2 way ANOVA with Sidak posthoc analysis comparing only YF-YM, YF-AF, YM-AM, AF-AM, adjusted p = *0.05, **0.01, ***0.001, ****<0.0001.

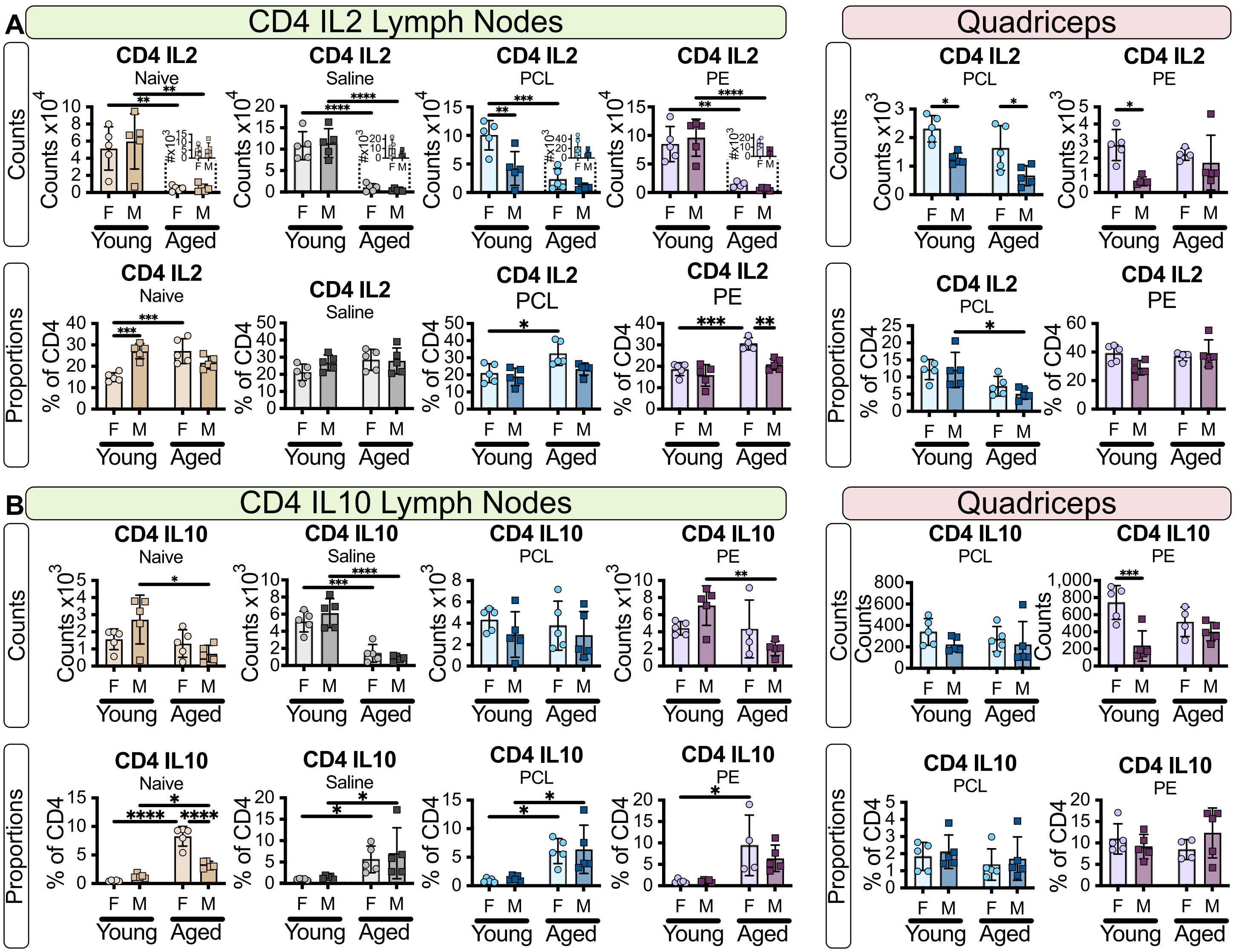


Supplemental Figure 15: (A) Counts and proportions for IL2 expression in CD4 T cells form the iLN and quadriceps. (B) Counts and proportions of IL10 in CD4 T cells in the iLN and quadriceps. Statistics: 2 way ANOVA with Sidak posthoc analysis comparing only YF-YM, YF-AF, YM-AM, AF-AM, adjusted p = *0.05, **0.01, ***0.001, ****<0.0001.

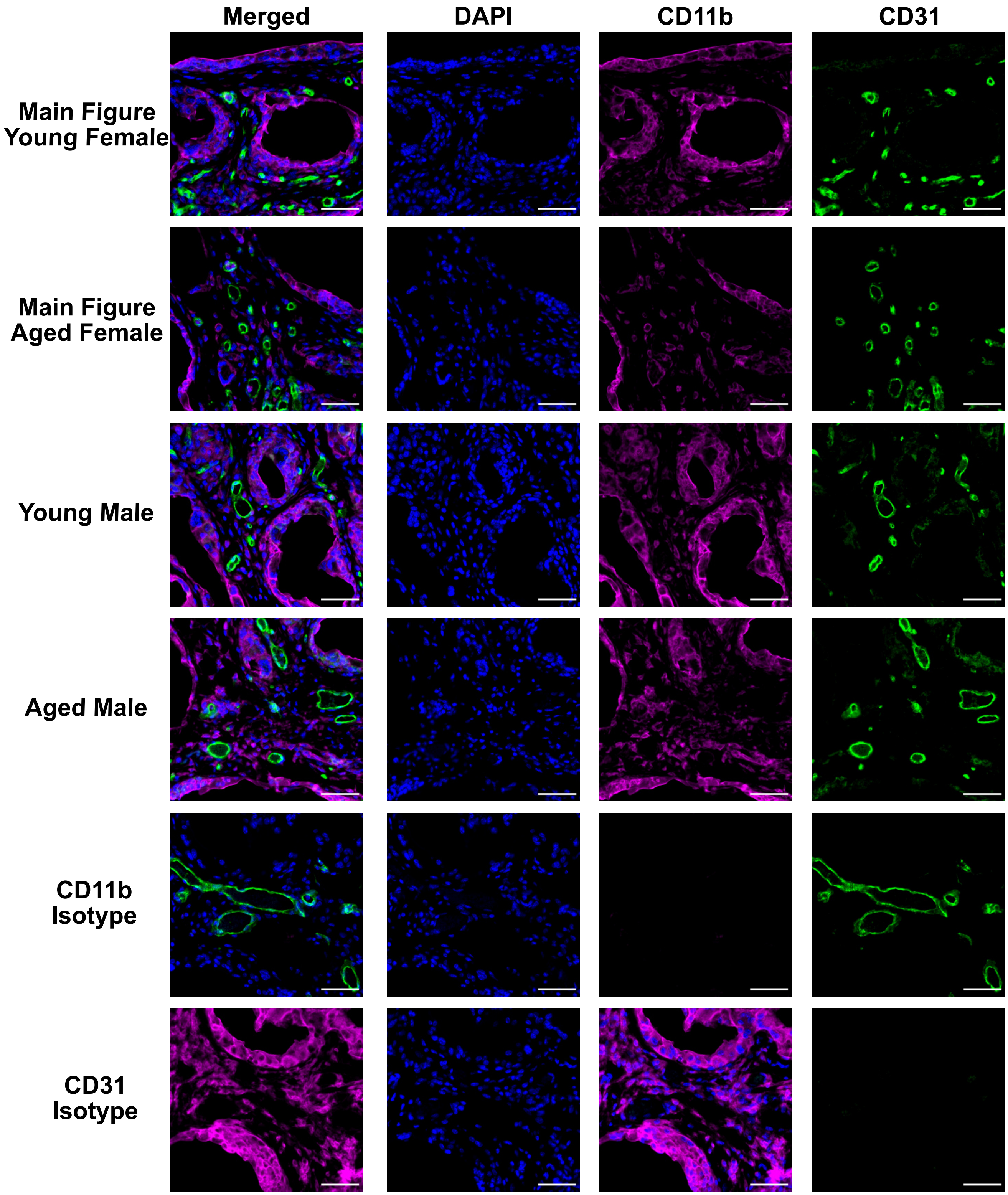


Supplemental Figure 16: Representative images for both age and sexes with corresponding single channel images and isotype controls for Figure 3A. Scale bars = 50 µm. All images acquired on a Zeiss Axio Imager A2 and linearly contrasted identically.

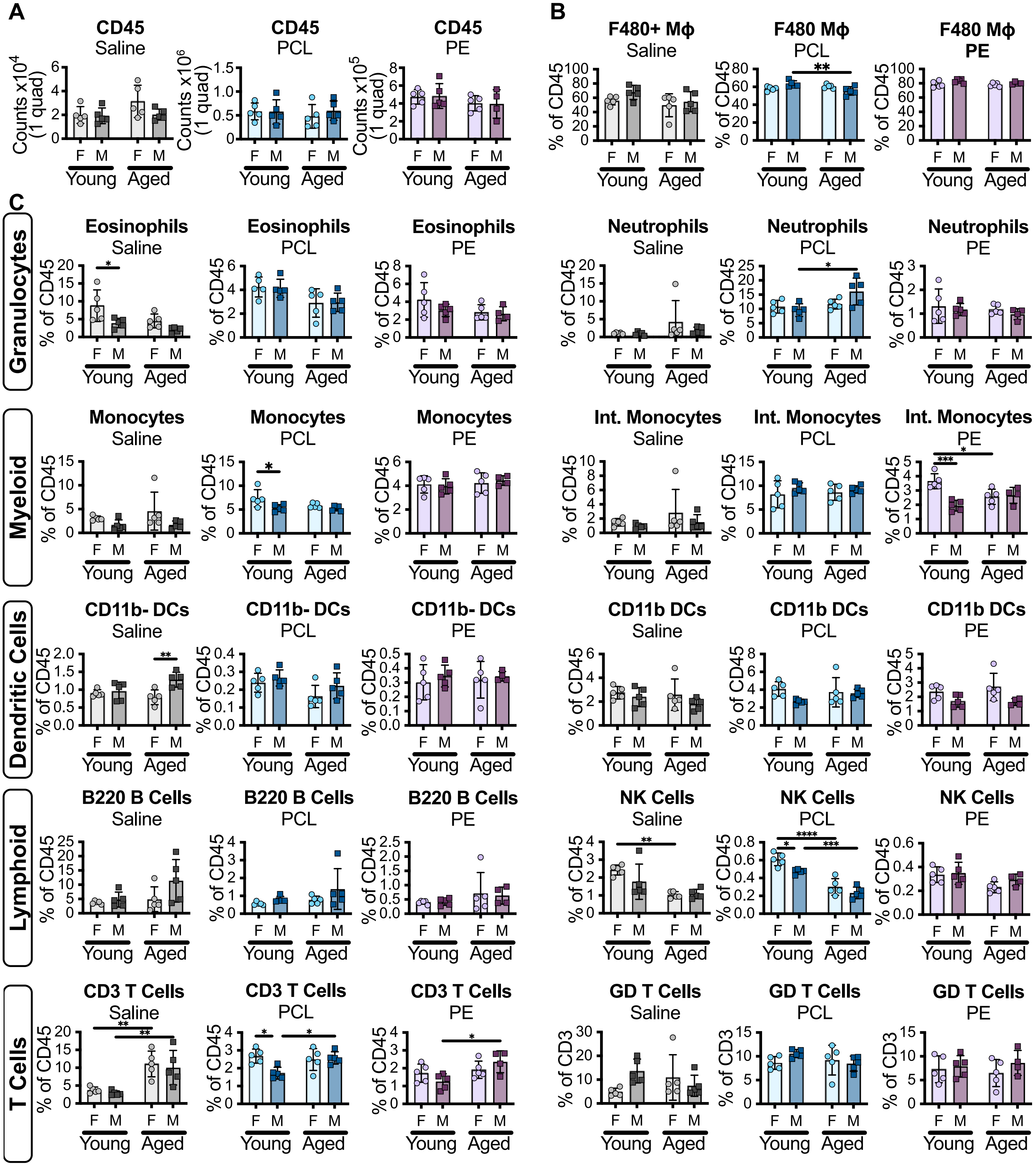


Supplemental Figure 17: (A) Calculated cell counts for 6-week VML- saline, -PCL, and -PE for one quadricep with fibrosis processed by digestion for the pan-immune flow cytometry panel. (B) Proportion of macrophages (C) Proportions of major cell populations identified by flow cytometry showing condition specific changes for some populations. Statistics: 2 way ANOVA with Sidak posthoc analysis comparing only YF-YM, YF-AF, YM-AM, AF-AM, adjusted p = *0.05, **0.01, ***0.001, ****<0.0001.

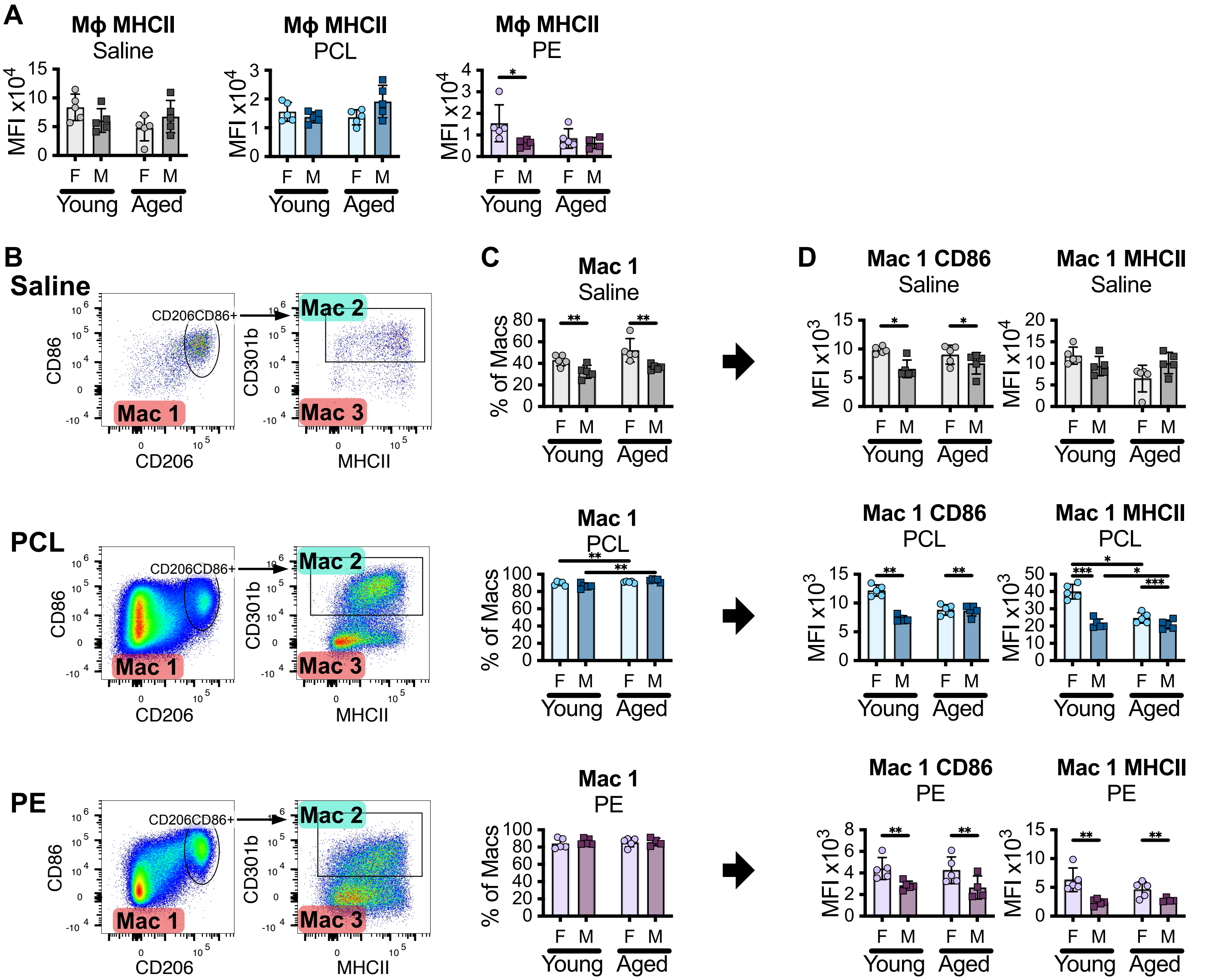


Supplemental Figure 18: (A) MHCII median fluorescence intensity for the F4/80 Macrophage population as a whole (B) Gating schematic for subpopulations of macrophages; a gate is set on the CD206 CD86 population and an identical not gate (red label) is used to identify Mac 1. Mac 2 is then gated from CD206 CD86 CD301b+ and Mac 3 is the CD301b not gate (C) proportions of the larger macrophage population in the particulate-induced fibrosis and the median fluorescence intensity for markers CD86 and MHCII. Statistics: 2 way ANOVA with Sidak posthoc analysis comparing only YF-YM, YF-AF, YM-AM, AF-AM, adjusted p = *0.05, **0.01, ***0.001, ****<0.0001.

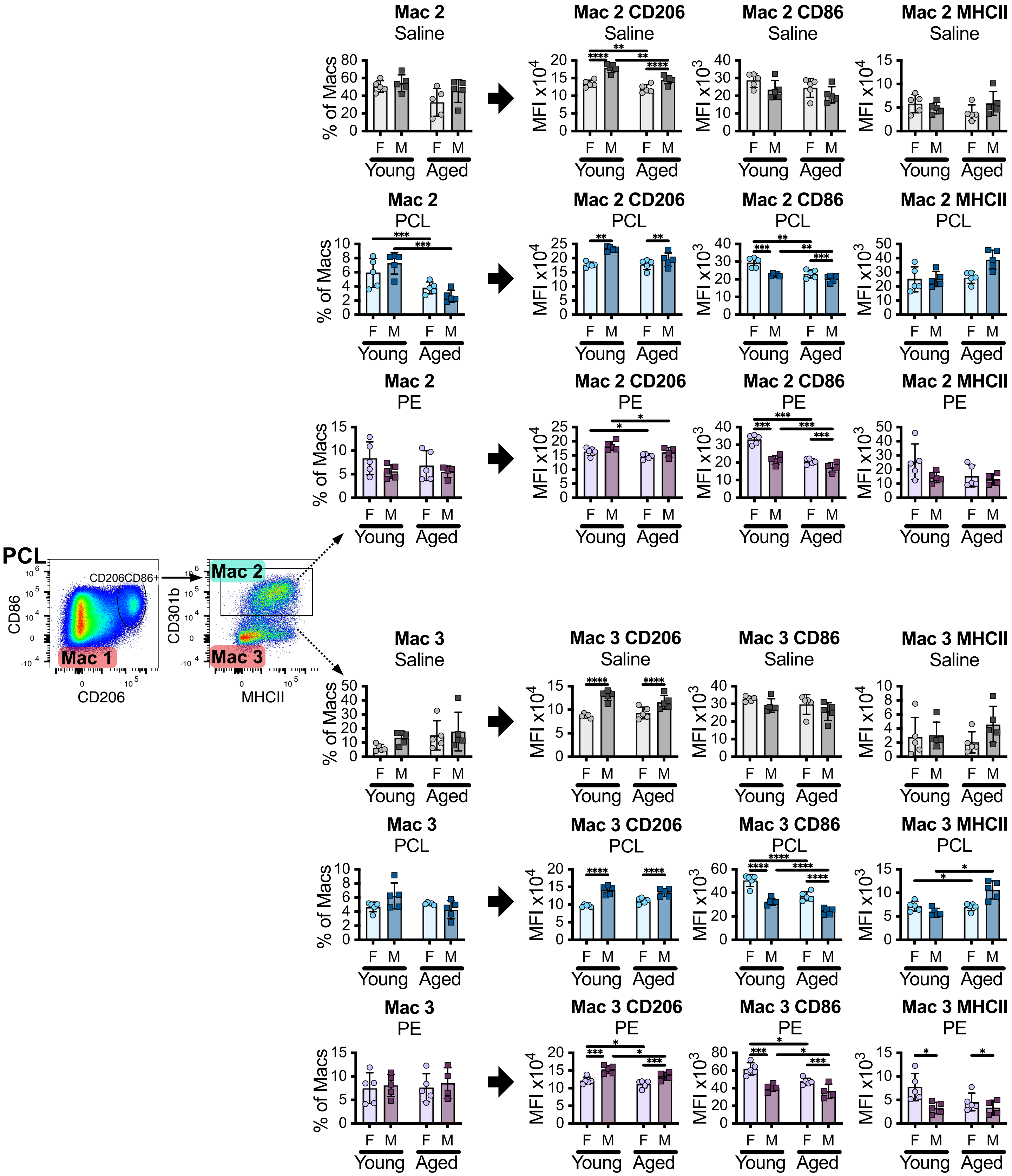


Supplemental Figure 19: Mac 2 and Mac 3 population proportions and the median fluorescence intensity for markers CD206, CD86, and MHCII. Statistics: 2 way ANOVA with Sidak posthoc analysis comparing only YF-YM, YF-AF, YM-AM, AF-AM, adjusted p = *0.05, **0.01, ***0.001, ****<0.0001.

Supplemental Table 3: Immunofluorescence Staining Primary and Secondary

| Antigen | Clone | Host Species | Stock Concentration (mg/mL) | Dilution | Manufacturer | Cat # | RRID |
|---|---|---|---|---|---|---|---|
| CD31 | EPR17259 | Rabbit | 0.519, 0.568 | 2000 | Abcam | ab182981 | AB_2920881 |
| p16 | EPR20418 | Rabbit | 0.583 | 1000 | Abcam | ab211542 | AB_2891084 |
| CD8a | EPR21769 | Rabbit | 0.526 | 1000 | Abcam | ab217344 | AB_2890649 |
| CD11b | EPR1344 | Rabbit | 1.266 | 1000 | Abcam | ab133357 | AB_2650514 |
| CD4 | EPR19514 | Rabbit | 0.687 | 1000 | Abcam | ab183685 | AB_2686817 |
| IgG (Isotype) | EPR25A | Rabbit | 1.675 | Match to primary dilution | Abcam | ab172730 | AB_2687931 |
| DAPI | – | – | – | 10 | Akoya | FP1490 | not available |
| Opal 520 | – | – | – | 100 | Akoya | FP1487001KT | not available |
| Opal 570 | – | – | – | 150 | Akoya | FP1488001KT | not available |
| Opal 650 | – | – | – | 150 | Akoya | FP1496001KT | not available |
